# Exposure to a mixture of organic pollutants in a threatened freshwater turtle *Emys orbicularis*: effects of age, sex, and temporal variation

**DOI:** 10.1101/2025.06.29.658864

**Authors:** Leslie-Anne Merleau, Olivier Lourdais, Anthony Olivier, Marion Vittecoq, Fabrice Alliot, Sira Traore, Carole Leray, Aurélie Goutte

## Abstract

Freshwater ecosystems constitute major sinks for organic contaminants, increasing anthropogenic pressures and threatening the unique biodiversity they harbour. In addition to persistent legacy compounds, such as polychlorinated biphenyls (PCBs) and organochlorine pesticides (OCPs), various pollutants are less persistent but are chronically released, including polycyclic aromatic hydrocarbons (PAHs), phthalate diesters (PAEs), pyrethroid pesticides, and insect repellent. Heretofore, these pollutants have received insufficient attention in freshwater reptiles, considering their potential to trigger detrimental effects on organisms. During two years (2019 and 2020), we quantified plasma levels of 46 compounds from 7 chemical families in two monitored populations of the protected European pond turtle (*Emys orbicularis*) in the Camargue wetland, France. PAHs and PAEs were found predominantly and concomitantly, with similar occurrences and levels in the two populations. We observed similar inter-annual variations in PAHs and PAEs with differences between males and females, highlighting the need for a better assessment of the role of sex in the exposure pathway and the toxicokinetics of contaminants, especially in turtles. The negative relationship between PAH levels and age, as well as the high intra-individual variation in levels of both contaminant families, provides further evidence of limited bioaccumulation of these pollutants in the blood of *E. orbicularis*. This could be explained by the metabolic biotransformation of parent compounds, highlighting the need to quantify the levels of PAH metabolites and phthalate monoesters. Finally, our work underscores the importance of long-term monitoring to better determine the vulnerability of turtle populations already exposed to a wide range of contaminants.

## Introduction

Freshwater ecosystems are continuously exposed to a wide range of organic pollutants from agricultural, urban, and industrial sources. These include legacy contaminants such as organochlorine pesticides (OCPs) and polychlorinated biphenyls (PCBs), as well as currently emitted compounds like polycyclic aromatic hydrocarbons (PAHs), phthalate esters (PAEs), and pesticides (Malaj et al., 2014; Rasmussen et al., 2015). Most Persistent Organic Pollutants (POPs) are banned by international regulations, such as PCBs and OCPs, but their persistence in aquatic environments still poses a long-term threat to biodiversity (Lohmann et al., 2007; Mateo et al., 2016). Some PAHs, and to a lesser extent PAEs, can also persist, notably in sediment (Manzetti, 2013; Net et al., 2015). However, international regulations targeting industrial and domestic releases of PAHs and PAEs remain limited or nonexistent, leading to steady environmental contamination (Ravindra et al., 2008; Malaj et al., 2014; Net et al., 2015).

Once in the environment, some of these contaminants can accumulate in aquatic organisms. PCBs and OCPs tend to biomagnify through food webs because of their lipophilicity and slow metabolism (Mackintosh et al., 2004; Walters et al., 2016; Goutte et al., 2020), whereas PAHs and PAEs are more prone to trophic dilution due to higher metabolization and excretion in vertebrate species (Mackintosh et al., 2004; Wan et al., 2007; Walters et al., 2016; Goutte et al., 2020; Lorrain-Soligon et al., 2025). However, repeated contamination through water and sediment can lead to chronic exposure and the saturation of metabolic capacities for these molecules (Van Der Oost et al., 2003; Tyler et al., 2018). For species with pulmonary respiration, ambient air pollution is another source of contamination (Teil et al., 2016).

Exposure to such contaminants can lead to detrimental effects in a wide range of organisms. PCBs, OCPs, PAHs, and PAEs can cause endocrine disruption, particularly in the thyroid and reproductive hormonal systems (Vethaak & Legler, 2013; Matthiessen et al., 2018; Wallace et al., 2020). Impairment of immunological functions, oxidative stress, genotoxicity, and embryotoxicity have also been reported in vertebrate species exposed to these contaminants (Reynaud & Deschaux, 2006; Marasco & Costantini, 2016; González-Mille et al., 2019; Wallace et al., 2020). These various sublethal effects can ultimately compromise individual fitness through higher mortality rates or reduced reproductive outputs, potentially leading to population decline (Aulsebrook et al., 2020; Campioni et al., 2024). However, the majority of ecotoxicological studies on organic contaminants still focus on abiotic environmental monitoring, excluding wildlife exposure (Simms et al., 2025).

This knowledge gap is particularly concerning given the current state of freshwater ecosystems. Wetlands are among the most threatened ecosystems worldwide, and pollution constitutes a major issue (Malaj et al., 2014; Dudgeon, 2019; Reid et al., 2019). Vertebrate populations in wetlands are declining faster and sharper than in any other ecosystem (Reid et al., 2019), highlighting the necessity for targeted monitoring of contaminant burdens in wildlife species (Simms et al., 2025). Furthermore, field studies have frequently focused on a specific class of pollutants, thereby neglecting the diversity of contaminants and potential synergistic effects (Relyea, 2009; Molbert et al., 2019).

The distribution of field studies across taxa is also uneven. Among vertebrates, reptiles remain largely overlooked in ecotoxicology despite being one of the most threatened taxa (Sparling et al., 2010; Simms et al., 2025). Nevertheless, some organic contaminants have been quantified in wild reptile species (Yu et al., 2012; Meyer et al., 2016; González-Mille et al., 2019; Sanjuan et al., 2023). Freshwater turtles are key species for monitoring pollutant exposure because of their high trophic positions and slow generational turnover that increase their susceptibility to bioaccumulation (Rowe, 2008). In addition, their lifespans and high site fidelity (Olivier, 2002; Fay et al., 2023) enable them to be exposed to pollutants over decades, providing insights into local contamination patterns (Rowe, 2008; Merleau et al., 2024a). The European pond turtle (*Emys orbicularis*) has a large Eurasian distribution with fragmented and declining populations, exposed to a wide range of anthropogenic disturbances. Several studies have investigated levels of metallic trace elements (MTEs), pesticides, and POPs, suggesting that this species is a good candidate for ecotoxicological studies (Swartz et al., 2003; Namroodi et al., 2017; Guillot et al., 2018; Beau et al., 2019; Burkart et al., 2021; Merleau et al., 2024a, 2024b).

We measured organic contaminants in two neighboring populations of European pond turtles monitored since 1976 (Olivier, 2002; Olivier et al., 2010) in the Camargue wetland (Southern France), a biodiversity hotspot. These two populations are located in the Rhône Delta and exposed to several sources of urban and industrial pollution (Roche et al., 2000; Berny et al., 2002; Ribeiro et al., 2005; Burkart et al., 2021). The Rhône River, one of France’s main catchment areas, is particularly exposed to PCBs and PAEs (Paluselli et al., 2018; Dendievel et al., 2020), and a major industrial port zone (ZIP) and petrochemical complex are located a few kilometers downstream. The region is also characterized by intensive rice cultivation, which is historically associated with heavy OCP use (Roche et al., 2000). The habitats of the two populations of *E. orbicularis* differ in their hydrology in relation to the irrigation and drainage systems of the rice fields. In the Camargue, rice paddies could regulate the intakes of PCBs and OCPs, that originate from the Rhône River, in the hydrosystem (Roche et al., 2009). Recent works have shown contrasting exposure to currently-used pesticides, metallic trace elements, and POPs in these two populations (Burkart et al., 2021; Merleau et al., 2024a, 2024b). Furthermore, our long-term monitoring provides insight into the effects of individual characteristics on contamination, such as age and sex. Using a multiresidue method of quantification adapted for plasma matrix (Molbert et al., 2019), we measured plasma concentration of 15 PAHs, 7 PAEs, and 24 other organic contaminants (PCBs, OCPs, pyrethroids, DEET, and PBDEs) over two years (2019 and 2020) in 162 individuals. We aimed to assess the spatiotemporal factors that influenced organic contaminant burdens in *E. orbicularis* in relation to chemical family; the individual characteristics such as sex, body condition, and age that shaped the levels of contaminants; and contaminant fluctuations at the intra-individual level in a subset of individuals sampled over the two years.

## Material and methods

### Animal captures and marking

Turtles were captured in two populations located in the Camargue wetland (Southern France) in the Natural Reserve of the Tour du Valat (43°30′N, 4°40′E; Fig. 1). These two populations of *E. orbicularis* are only a few kilometers apart, but their habitats differ strongly regarding the hydrological characteristics (Olivier et al., 2010; Burkart et al., 2021; Merleau et al., 2024a, 2024b). Estimated population sizes are presented in Table 1 (Ficheux et al., 2014). The population of the Esquineau occupies canals and marshes associated with the rice paddies irrigation system, using water coming from the Rhône River (Fig. 1). Previous work on the same individuals showed that turtles from this population had higher levels of blood mercury and lead compared to the population of Faïsses (Merleau et al. 2024a). This second population is found in the paddy drainage system and the associated canals and marshes (Fig. 1) and presents higher occurrences and levels of pesticides used in the rice fields (Merleau et al. 2024b).

**Table 1.** Estimated population sizes and distribution of sample group sizes by population, year, and sex.

| Population | Estimated Size<br>(Ficheux et al. 2014) | Year | N<sub>Tot</sub> Female | N<sub>KA</sub> Female | N<sub>Tot</sub> Male | N<sub>KA</sub> Male |
| --- | --- | --- | --- | --- | --- | --- |
| Esquineau | 300 ± 90 | 2019 | 34 | 22 | 24 | 19 |
|  |  | 2020 | 37 | 24 | 24 | 12 |
| Faïsses | 73 ± 21 | 2019 | 29 | 8 | 20 | 6 |
|  |  | 2020 | 34 | 14 | 16 | 6 |

Note: N<sub>Tot</sub>: number of samples for the whole set of sampled individuals; N<sub>KA</sub>: number of samples from individuals of known age

**Figure 1.**
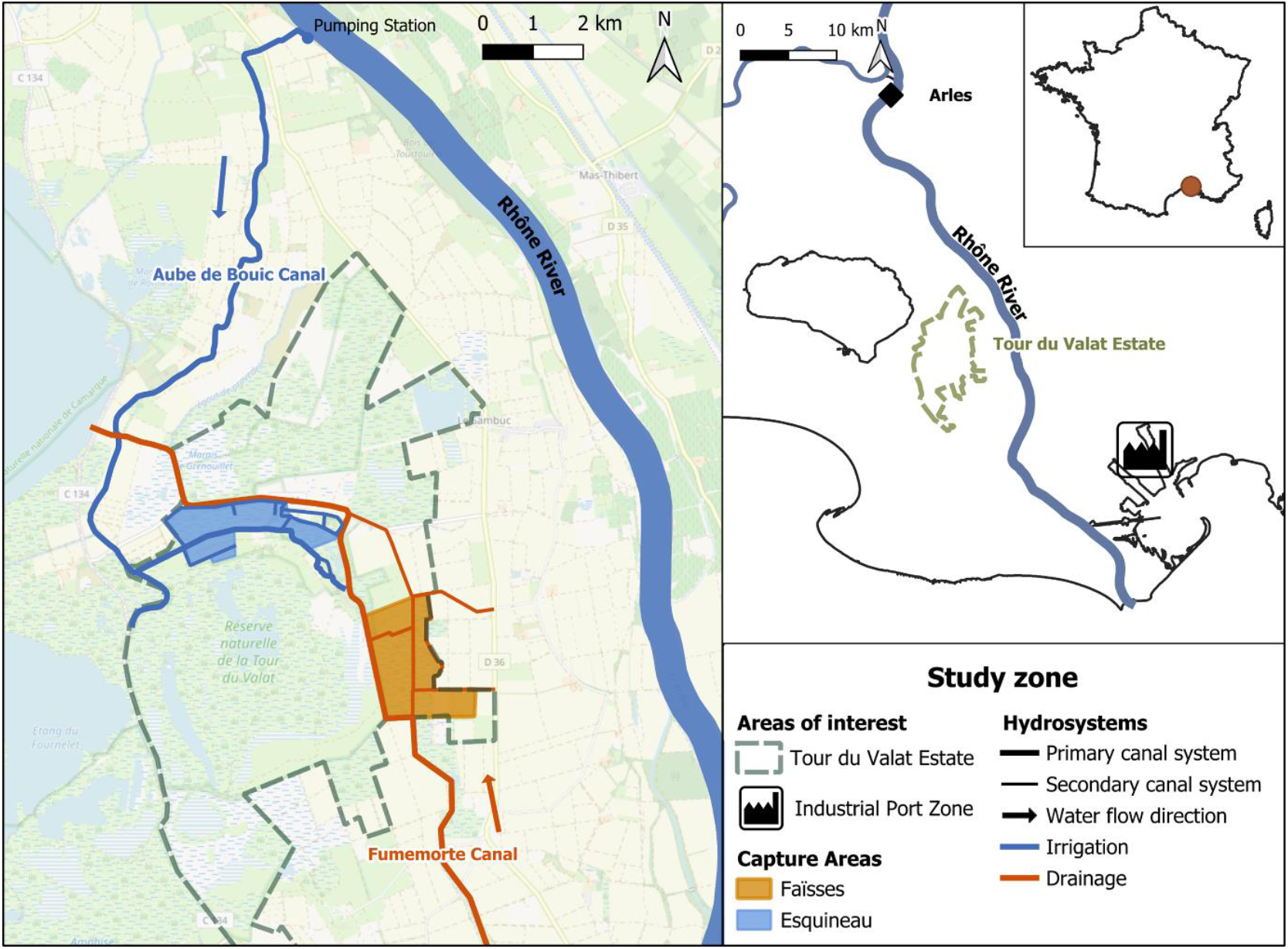
Study zone and capture areas of two populations of *E. orbicularis* in the Camargue wetland (France), in relation to the hydrosystems and the anthropic activity zones (blue area: Esquineau, irrigation canals and marshes, orange: Faïsses, drain canals and marshes, green dashed polygon: Tour du Valat Estate).

Turtles were collected from the 23rd of April to the 26th of July in 2019 and from the 9th of April to the 7th of August in 2020, either manually or with funnel traps. Turtles are individually recognized by notching their marginal and nuchal scales as part of a long-term capture-mark-recapture survey (1976–2025) (Olivier, 2002; Ficheux et al., 2014). Traps located in 20 and 16 capture stations (Esquineau and Faïsses, respectively) were checked every day, and individuals were released at the precise location of capture. Captures of this protected species were performed under French Departmental Authorities authorization (Permits: DREAL Cerfa 13616-01; N°13-2020-03-27-007).

### Biometrics and blood sampling

Turtles were weighed with a precision scale (Mettler Toledo PB3001-S) to the nearest gram and measured with a caliper. We used the dorsal shell length (hereafter “carapace length”) as the measure of the size. We estimated body condition using the residuals of a linear regression of the log-transformed mass against the log-transformed carapace length with the addition of sex as a control variable. Thanks to the CMR monitoring, the age of the individual was known for 111 samples (68 from 48 females, range: 5-32 years; 43 from 35 males, range: 4-25 years; Table 1). When the first capture of an individual happens before 5 years old, the age is obtained by counting the number of growth rings on the plastron (Castanet, 1988; Olivier, 2002).

In order to comply with animal welfare standards, we performed blood sampling on turtles weighing more than 300 g, i.e., sampling volume corresponding to less than 1% body mass. When collecting and processing biological samples, we were careful to ensure that plastic sampling materials did not contain PAEs, and we did not use polyvinyl chloride (PVC) to avoid contamination of biological samples. We sampled 1.5 mL of blood from 162 individuals using the dorsal coccygeal vein and a previously heparinized syringe and a 25G needle. During the study, 56 individuals were sampled both years (2019 and 2020), resulting in a total of 218 blood samples. After centrifugation during three minutes at 3000 rpm to separate plasma from erythrocytes, the samples were kept at -18 °C until analyses at Sorbonne University’s UMR 7619 METIS in 2022. All samples in this study are part of a set of samples on which pesticide and trace element quantifications have been previously performed and published (Merleau et al., 2024a, 2024b). An Ethics Committee approved this procedure in accordance with EU Directive 2010/63/EU for animal experiments (permit: APAFIS 17899–201 812 022 345 423).

### Multiresidue analytic protocol

Using a method developed by Molbert et al., (2019), we quantified 15 PAHs (acenaphtylene, naphthalene, anthracene, fluorene, phenanthrene, fluoranthene, pyrene, chrysene, benzo[a]anthracene, benzo[a]pyrene, benzo[b]fluoranthene, benzo[k]fluoranthene, benzo[g,h,i]perylene, dibenzo[a,h]anthracene, indeno[1,2,3-cd]pyrene), 7 PAEs (dimethyle phthalate (DMP), diethyle phthalate (DEP), di-iso-butyl phthalate (DIBP), di-n-butyl phthalate (DNBP), n-butyl benzyl phthalate (BBP), di-2-ethylhexyl phthalate (DEHP), di-n-octyl phthalate (DnOP), 7 pyrethroids (bifenthrin, permethrin, phenothrin, cyfluthrin, cypermethrin, fenvarelate, deltamethrin), 4 OCPs (PeCB, HCB, lindane, p,p’-DDE), 7 PCB congeners (28, 52, 101, 118, 138, 153, 180), and 6 polybrominated diphenyl ethers (PBDE) congeners (28, 47, 99, 100, 153, 154). Quantification was carried out by internal calibration using a mix of standards (phenanthrene d10; pyrene d10; benzo(a)anthracene d12; DEP d4; DEHP d4; lindane-d6; cis-permethrin 13C; PCB 30/107, and BDEs ^13^C-BDE-47 and ^13^C-BDE153; Supplementary Material, Table S3) that were added to 100 µL of thawed plasma. After being vortexed and stored one night at 4 °C, the extraction was performed with hexane/dichloromethane/acetone (1:1:1, v/v/v) with two rounds of centrifugation, then concentrated under a nitrogen stream to approximately 0.5 mL. Purification was performed on Florisil cartridges (Supelco 1 g/6 mL, Sigma-Aldrich) using two cycles of elution with 2×5 mL of hexane/dichloromethane (1:1, v/v) and 2×5 mL of hexane/acetone (1:1, v/v). After concentration to 0.5 mL under a nitrogen stream and with the Genevac Concentrator EZ-2 (Biopharma Technologies), the extracted samples were filtered with nylon filter tubes (porosity 0.2 μm, VWR, Fontenay-sous-Bois) and rinsed twice with 70 µL of hexane. Filtrates were obtained by centrifugation (6000 rpm, 2 min) and rinsing (50 µL hexane) of the tubes.

The samples were then analysed by gas chromatography coupled with tandem mass spectrometry (GC-MS/MS) using an Agilent 7890 A gas chromatograph (GC) coupled to a 7000 B triple quadrupole mass spectrometer (MS/MS) (Agilent Technologies). Following Molbert et al. (2019) protocol, the targeted compounds were characterized in the gas phase using electron ionization (EI, +70 eV) with an ionization source set at 250 °C. A Zebron SemiVolatile analytical column (30 m, 0.25 mm ID x 0.25 μm film thickness; Phenomenex) attached to a Restek deactivated silica guard column (0.25 mm ID) was used to separate molecules. A 1 μL sample extract was injected in splitless mode with the injector port temperature at 290 °C. The flow rates for the carrier gas (helium) and collision gas (nitrogen) were maintained at 1.6 mL/min and 1.5 mL/min, respectively. The oven temperature was programmed as follows: 4 min at 70 °C, then increased from 70 to 150 °C at 25 °C/min, then raised to 225°C at 3 °C/min and finally to 310 °C at 5 °C/min, before being held at 310°C for 10 min. The temperature of the MS/MS transfer line was set at 250 °C.

For each series of 12 samples, solvent blanks were prepared following the same procedure as the plasma samples. To correct for background contamination, blank correction was performed if plasma concentrations of target compounds were not four times higher than blank concentration (Laborie et al., 2016). The method was validated in terms of yield, repeatability, and quantification limits for these compounds and in this matrix (Supplementary Material, Tables S4, S5).

### Statistical analyses

We performed all statistical analyses with R software (4.4.2) and the packages lme4 (Bates et al., 2015), emmeans (Lenth, 2025), DHARMa (Hartig, 2024), and rptR (Stoffel et al., 2017). Non-significant variables (p > 0.05) were gradually eliminated from the models. The statistical values acquired in the final stage prior to their removal from the models are supplied for non-significant variables. Tukey post hoc tests in the emmeans package were used for pairwise comparisons (Lenth, 2025). Using the DHARMa package, residual analysis was performed to check model assumptions and to test outliers (Bates et al., 2015).

#### Variation in contaminants

To account for repeated sampling of individuals, we employed LMMs (Linear Mixed Models) with individual identity as a random effect. We examined how the sums of plasmatic concentrations of compounds from the same chemical family were affected by year, month, site, sex, carapace length, body condition, and the interactions site × year, sex × year, sex × size, and sex × age. To best suit the model assumptions, the sums of PAEs and PAHs were log-transformed. We also performed the same LMMs by examining the impact of age on the subset of known-age individuals (n = 83), removing size and body condition variables from the models as they were highly associated with age.

#### Intra-individual variation over time

We also assessed annual variations at the intra-individual level using repeated measures from 56 individuals sampled in 2019 and 2020. We used t-tests to compare groups of individuals sampled each year in order to evaluate variations within individuals collected across the two years. The differences were tested for normality using the Shapiro-Wilk normality test. For the PAH concentrations, as the Shapiro-Wilk test indicated a statistically significant deviation from normality, we confirmed the results of the t-test using a Wilcoxon test, which confirmed the statistical result obtained. Using the program rptR, we then conducted individual repeatability tests to evaluate the intra-individual variation, i.e., the intraclass correlation coefficients (ICCs) or repeatability index (R), of contaminant concentrations over time between 2019 and 2020 (Stoffel et al., 2017). ICCs are calculated by dividing the inter-individual variance by the sum of inter- and intra-individual variances. A repeatability index < 0.4 indicates poor reproducibility within individuals, while indicating excellent reproducibility when > 0.75 (Rosner, 2010). “Repeatability” is defined as the stability of the intra-individual measured contaminants over time.

## Results

### Contaminant levels in the plasma of *E. orbicularis*

Organic pollutants were detected in 93.1% of the plasma samples. On average, 3 different chemicals were quantified per sample, and only 15.1% of the samples contained a single compound. The highest number of compounds detected in a single sample was 11, observed in one individual, and 58.3% of the samples contained compounds from at least 2 chemical families.

PAEs were the compounds with the highest quantification rates (78.4%), followed by the PAHs (66.5%). Among PAEs, DIBP and DNPB were the most frequently quantified (49.1% and 43.1%, respectively) and had the maximum mean concentrations, 380.9 and 187.4 ng/mL of plasma, respectively. Despite being quantified in substantial proportions, DEP, DEHP, and DMP had lower plasma concentrations (41.4, 32.6, and 4.8 ng/mL, respectively). BBP and DnOP were below the LOQ in all the samples.

Regarding PAHs, naphthalene was the most quantified (54.1%) with the highest mean plasmatic concentration (5.2 ng/mL). Two other PAHs were quantified in substantial frequencies: phenanthrene (23.9%) and fluorene (22.9%). However, their plasmatic levels were lower than naphthalene: 3.4 and 0.7 ng/mL, respectively. Fluoranthene, pyrene, dibenzo[a,h]anthracene, indeno[1,2,3-cd]pyrene, and benzo[g,h,i]perylene were quantified in less than five samples and thus were not included in the sum of PAH concentrations. The summary of PAE and PAH plasmatic concentrations, by site and by year, is available in Supplementary Material (Tables S1, S2).

We quantified DEET in 19 samples (8.7%). Among the PCB congeners, we quantified PCB 28, 52, and 138 in 3, 10, and 1 samples, respectively. No pyrethroids, OCPs, or PBDEs were quantified.

#### PAE concentrations

The sum of the PAE concentrations differed between males and females in interaction with the year (Table 2). Males had significantly higher levels compared to females, but only in 2020 (Fig. 2). We did not detect a significant effect of the month of capture (Table 2) nor a difference between the two populations when all the samples were taken into account (Table 2). Regarding individual characteristics, plasmatic levels of PAEs were not affected by individual size and body condition (Table 2). Although the final model for the subset of known-age individuals included the site of capture as an explanatory variable, this effect was nearly non-significant (Table 2). We also found an effect of the year: plasmatic levels of PAEs were lower in 2019 than in 2020 (Table 2, Fig. 2). The levels of PAEs significantly decreased with age (Table 2, Fig. 3).

**Table 2.** Effects of environmental and individual variables on the levels of PAEs and PAHs in the plasma of *E. orbicularis* in two populations of the Camargue wetland (France). Models were selected using a top-to-bottom approach, based on two general models: log(Σcontaminants+1) ∼ population*year + month + sex*year + sex*scale(CL) + body condition and log(Σcontaminants+1) ∼ population*year + sex*year + sex*age + month for the subset of known-age individuals. For each effect tested, we present either (1) if selected in the final model, the observed significant effect (in bold), or (2) if removed, the level before exclusion.

|  | All (218 samples) |  |  | Known-age (111 samples) |  |  |
| --- | --- | --- | --- | --- | --- | --- |
| A. Sum of 5 PAEs | chi<sup>2</sup> | df | p value | chi<sup>2</sup> | df | p value |
| Population*Year | 0.525 | 1 | 0.469 | 1.702 | 1 | 0.192 |
| **Year** | **7.739** | **1** | **0.005** | **22.191** | **1** | **< 0.001** |
| **Population** | 0.020 | 1 | 0.889 | **4.922** | **1** | **0.027** |
| Month | 1.070 | 4 | 0.899 | 5.375 | 4 | 0.251 |
| **Sex\*Year** | **5.300** | **1** | **0.021** | 1.683 | 1 | 0.858 |
| Sex*CL (scaled) | 0.275 | 1 | 0.6 | - | - | - |
| Sex*Age | - | - | - | 0.002 | 1 | 0.966 |
| Sex | 0.012 | 1 | 0.912 | 0.345 | 1 | 0.557 |
| CL (scaled) | 0.020 | 1 | 0.887 | - | - | - |
| Body condition (scaled) | 0.115 | 1 | 0.734 | - | - | - |
| **Age** | - | - | - | **7.765** | **1** | **0.005** |
| B. Sum of 3 PAHs |  |  |  |  |  |  |
| Population*Year | 0.040 | 1 | 0.841 | 0.041 | 1 | 0.840 |
| **Year** | **19.084** | **1** | **< 0.001** | **39.081** | **1** | **< 0.001** |
| Population | 0.688 | 1 | 0.407 | 0.128 | 1 | 0.721 |
| **Month** | **12.698** | **4** | **0.013** | 8.936 | 4 | 0.063 |
| Sex*Year | 0.156 | 1 | 0.693 | 0.047 | 1 | 0.828 |
| Sex*CL (scaled) | 1.420 | 1 | 0.233 | - | - | - |
| **Sex\*Age** | - | - | - | **11.912** | **1** | **0.001** |
| Sex | 0.072 | 1 | 0.788 | 0.607 | 1 | 0.436 |
| CL (scaled) | 0.437 | 1 | 0.509 | - | - | - |
| Body condition (scaled) | 0.653 | 1 | 0.419 | - | - | - |
| Age | - | - | - | 0.626 | 1 | 0.429 |

Note: Results of linear mixed models (LMMs) for the sums of PAE (phthalate diester) and PAH (polycyclic aromatic hydrocarbon) levels. Known-age: a subset of individuals of known age. CL: carapace length, a measure of the body size. The significant variables retained in the final model are in bold font. Sum of 5 PAEs: DIBP, DNPB, DEP, DEHP, and DMP. Sum of 3 PAHs: naphthalene, phenanthrene, and fluorene.

**Figure 2.**
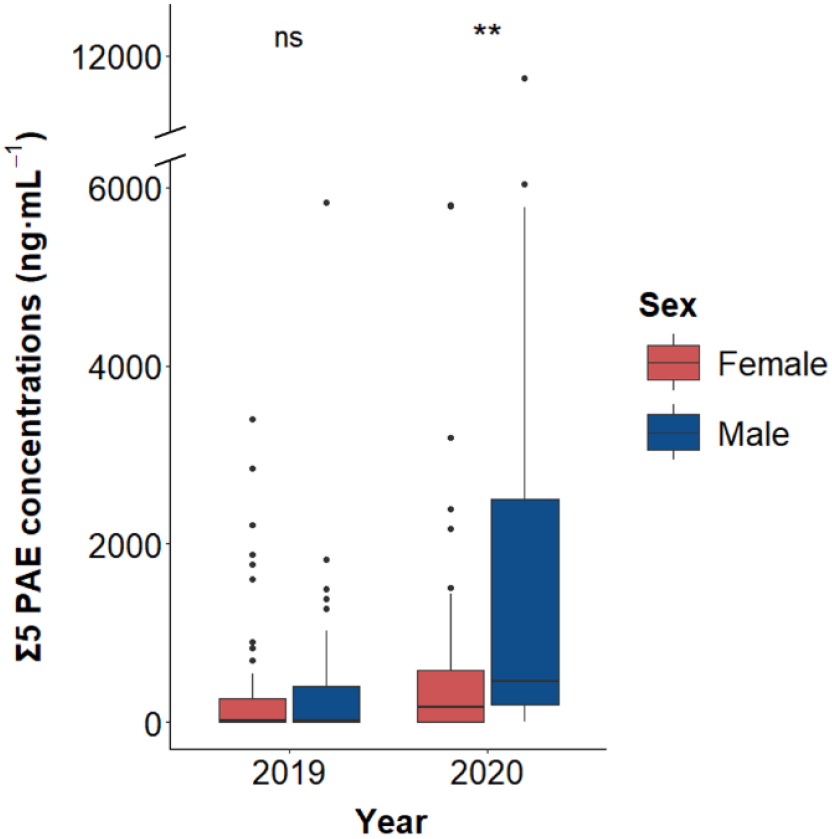
Sum (Σ) of the five phthalate diester (PAE) concentrations (ng.mL^-1^) quantified in the plasma of *E. orbicularis* from two populations in the Camargue (France) sampled in 2019 and 2020, according to the year and the sex of individuals (female: red; male: blue; n_Female_ = 134, n_Male_ = 84). Statistical significance obtained from the LMM model: **: p < 0.01, ns: not significant.

**Figure 3.**
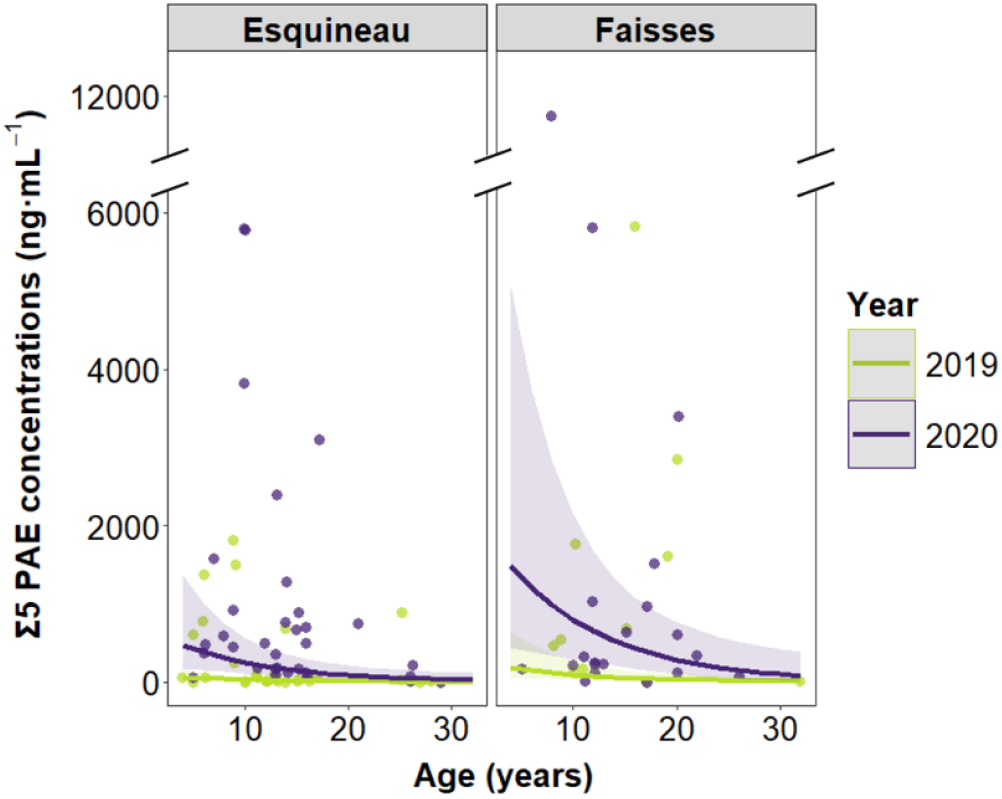
Sum (Σ) of the five phthalate diester (PAE) concentrations (ng.mL^-1^) quantified in the plasma of *E. orbicularis* from the Camargue (France) sampled in 2019 and 2020, according to the age of the individuals, the year (2019: green; 2020: purple), and the population (dots: data; lines: predictions of the LMM model including the age, the year, and the population; R^2^ conditional = 0.34, n_2019_ = 55, n_2020_ = 56).

#### PAH concentrations

The plasmatic levels of PAHs were lower in 2019 compared to 2020 (Table 2, Fig. 4) and were variable between months, although post-hoc tests did not show statistical differences between pairs of months (Fig. S1). We did not detect any other effect on the sum of PAH concentrations when considering all the samples. However, for samples from known-age individuals, we also detected an interaction between sex and age (Table 2). Older females had higher PAH levels, whereas this correlation was negative for males (Fig. 5). Females reached higher ages in our dataset, but the interaction between sex and age was still observed when retrieving older females. However, females’ levels were stable when only individuals under 25 years old were considered, indicating that levels increased later in females.

**Figure 4.**
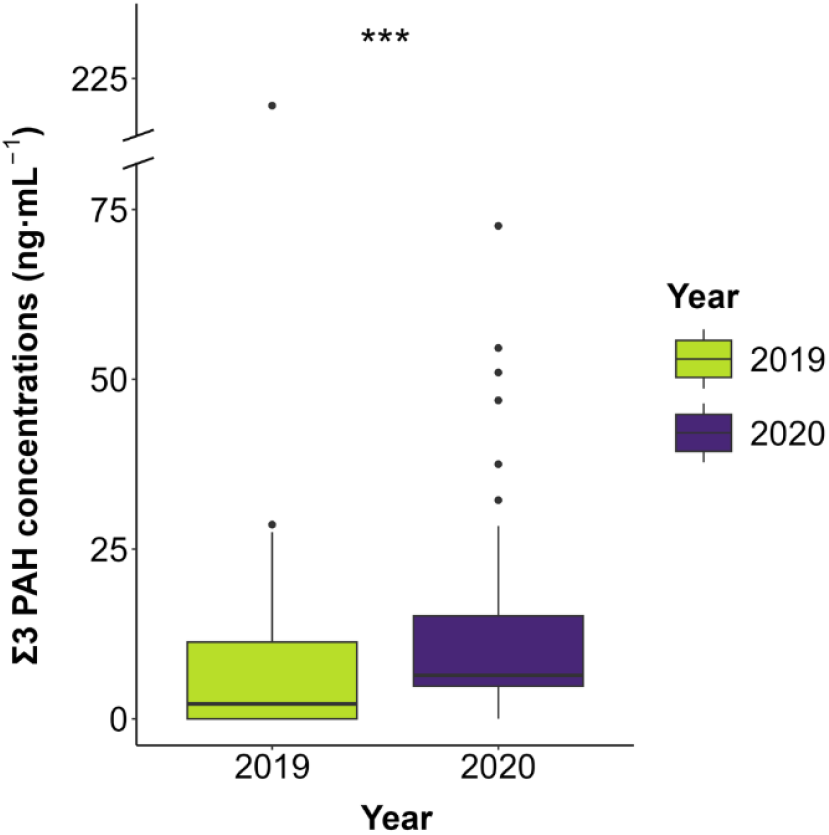
Sum (Σ) of the three polycyclic aromatic hydrocarbon (PAH) concentrations (ng.mL^-1^) quantified in the plasma of *E. orbicularis* from two populations in the Camargue (France) sampled in 2019 and 2020 (2019: green; 2020: purple; n_2019_ = 107; n_2020_ = 111). Statistical significance obtained from the LMM model: ***: p < 0.001.

**Figure 5.**
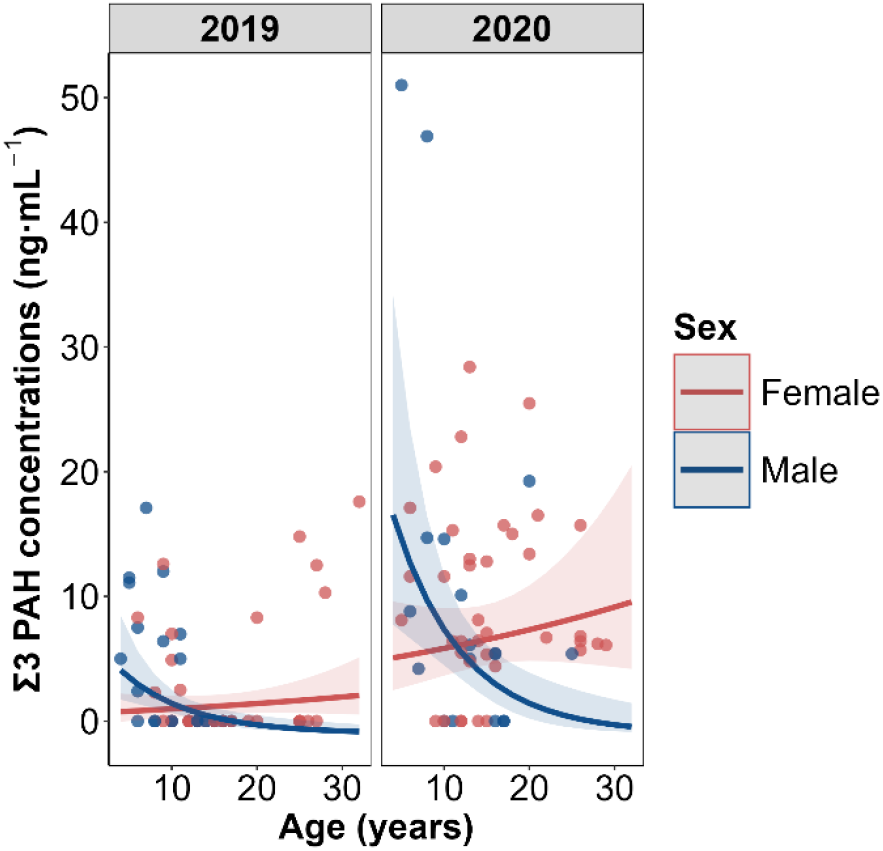
Sum (Σ) of the three polycyclic aromatic hydrocarbon (PAH) concentrations (ng.mL^-1^) quantified in the plasma of *E. orbicularis* from the Camargue (France) sampled in 2019 and 2020, according to the age of the individuals, the sex (female: red; male: blue), and the year (dots: data; lines: predictions of the LMM model including the age, the sex, and the year; R^2^ conditional = 0.38; n_Female_ = 68; n_Male_ = 43).

### Intra-individual variation

Using the contamination data of the 56 individuals sampled during two years, we were able to assess the inter-annual variation at the individual level. Individuals exhibited significantly higher levels of PAEs and PAHs in 2020 compared to 2019 (paired t-tests: PAEs, t = -3.017, df = 55, p = 0.004; PAHs, t = -4.047, df = 55, p < 0.001). We found low and non-significant intra-individual repeatability for both the PAE and the PAH concentrations, with repeatability coefficients of 0.065 and 0.108, respectively (Fig. 6).

**Figure 6.**
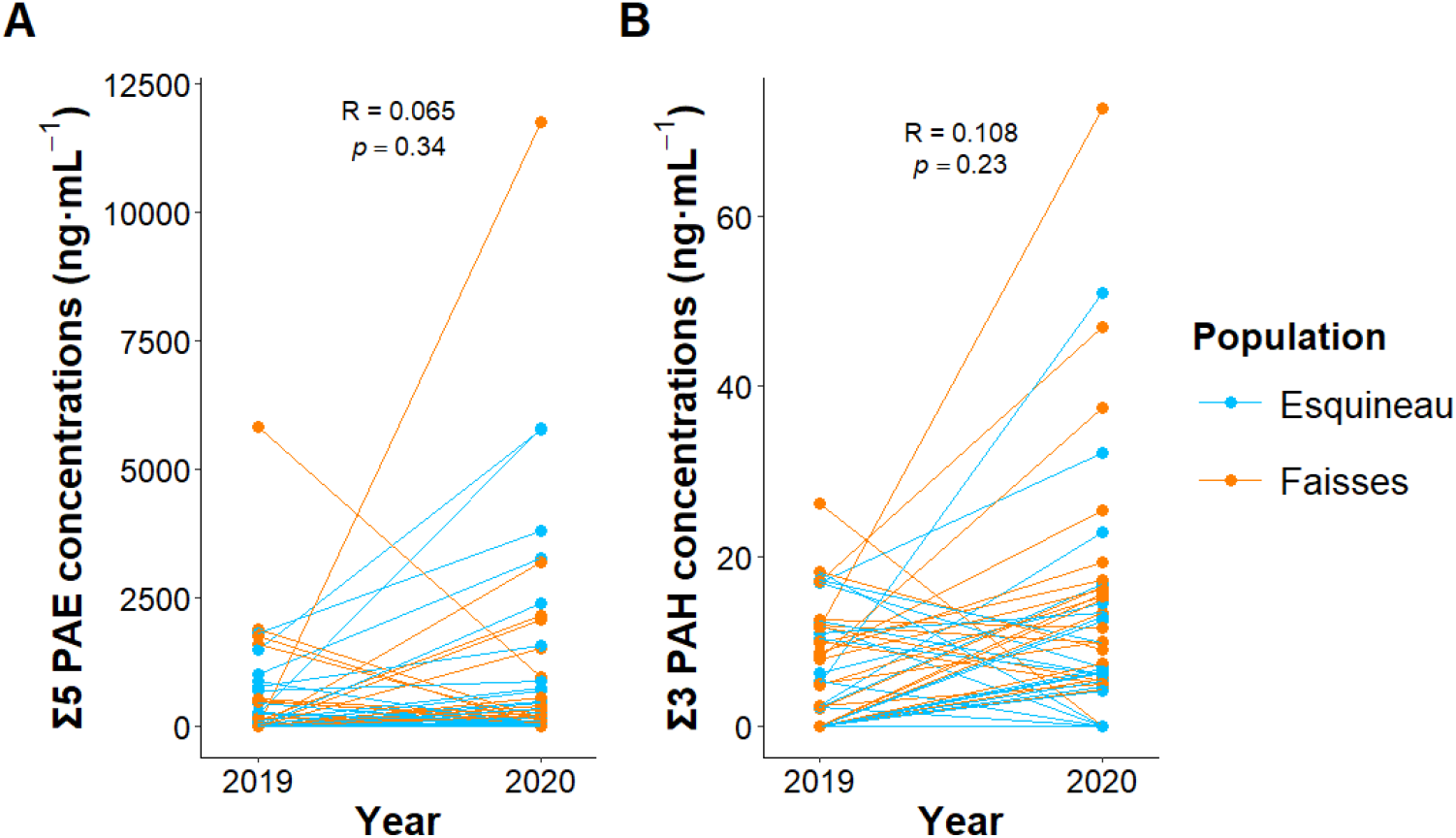
Sum (Σ) of (A) the five phthalate diester (PAE) and (B) the three polycyclic aromatic hydrocarbon (PAH) concentrations (ng.mL^-1^) quantified in the plasma of *E. orbicularis* from the Camargue (France) sampled both in 2019 and 2020, n = 56 (Esquineau: blue; Faïsses: orange; R: repeatability coefficient; p: significance).

## Discussion

In this study, we investigated the circulating levels of organic compounds from 7 major chemical families in two populations of *E. orbicularis*, a threatened freshwater turtle species, in relation to spatiotemporal characteristics and individual traits. PAHs and PAEs were quantified predominantly, whereas DEET and PCBs exhibited low detection frequencies, and pyrethroids, OCPs, and PBDEs were below the limits of quantification. At the spatiotemporal level, the year of sampling emerged as the strongest driver of contaminant concentrations. In contrast, individual traits had limited influence, with age and sex contributing primarily through interactions, either between them or with other variables.

### Diversity of contaminants

The contaminant plasmatic burden of the two populations was overwhelmingly due to PAEs and PAHs, and the majority of the individuals were contaminated by both of these chemical families. These results highlight the ubiquity of these two families in the environment in the vicinity of the petrochemical complex of Fos-sur-Mer.

PAHs are characterized by their number of benzene rings and can thus be classified as low (LMW) or high molecular weight (HMW) (Ravindra et al., 2008). Our results showed that the PAHs quantified in the plasma of *E. orbicularis* were almost exclusively two- or three-ringed: naphthalene, phenanthrene, and fluorene. These PAHs are known to be dominant in the surface water of the area (Guigue et al., 2014). As LMW PAHs, they exhibit higher water solubility, less sorption on dissolved carbon matter, and thus slower sedimentation than PAHs with 4 or more rings, and thus may be more bioavailable for aquatic organisms (Meador et al., 1995; Rabodonirina et al., 2015). In addition, HMW PAHs appear to be less absorbed and more metabolized in higher vertebrates (Perugini et al., 2007; Frapiccini et al., 2020), which could explain the poor detection in our populations. The nature of the matrix should also be considered: hydrophilic compounds are more likely to be found in plasma than hydrophobic compounds. However, the lower yield rates of some HMW PAHs in our study might provide an additional hypothesis to explain the occurrence differences between HMW and LMW PAHs (Supplementary Material, Table S5). The high occurrence of naphthalene, phenanthrene, and fluorene was also observed in sea turtle *Caretta caretta* and *Chelonia mydas* populations (Camacho et al., 2012, 2013; Cocci et al., 2018; Sinaei & Zare, 2019; López-Berenguer et al., 2023). To our knowledge, circulating levels of PAHs in freshwater turtles were only observed in *Actinemys marmorata* populations in California (Meyer et al., 2016). The sums of the plasmatic PAHs in our populations were lower than those found in *A. marmorata*, mainly due to higher levels of pyrene in this species. In detail, phenanthrene concentrations were similar to our populations of *E. orbicularis*, but fluorene was two times higher (Meyer et al., 2016). Only one study has monitored PAH levels in *E. orbicularis* populations in Azerbaijan, quantifying mostly naphthalene and phenanthrene in fat tissues (Swartz et al., 2003), but the difference between the matrices limits the comparison with our results. However, a later study suggested that chromosomal damages in these populations were correlated with environmental levels of three-ring PAHs (Matson et al., 2005). In *C. caretta*, levels of circulating LMW PAHs have been highly correlated with increased DNA methylation in blood cells (Cocci et al., 2018). The genotoxicity of LMW PAHs has also been highlighted in both in vitro and in vivo studies in fish (Holth et al., 2009; Yazdani, 2020). Our results suggest that even though HMW-PAHs with high toxicity were not quantified, the LMW-PAH burden may have adverse effects on the two *E. orbicularis* populations.

We quantified 5 PAEs in the plasma of *E. orbicularis*, with higher occurrences for intermediate (DIBP, DNBP) and high (DEHP) molecular weight PAEs. These PAEs are the most abundant in freshwater environments and have various origins, from atmospheric deposition to leaching from plastic debris (Net et al., 2015). Despite the scarcity of PAE monitoring in wildlife, and in reptiles in particular, these 3 PAEs exhibit frequent quantifications in vertebrate populations (Savoca et al., 2018; Liu et al., 2019; Blasi et al., 2022; Molbert et al., 2025). To our knowledge, our work is the first to assess PAE burden in the plasma of a freshwater turtle species. In Mediterranean populations of *C. caretta*, plasma levels of DEHP were similar to our populations of *E. orbicularis*, whereas DIBP and DnBP were, respectively, 9 and 3 times lower (Blasi et al., 2022). PAEs do usually not induce mortality in vertebrates and mainly cause endocrine disruption (Oehlmann et al., 2009; Tyler et al., 2018), but long-term toxicity thresholds are unknown for reptile species. In mammal, fish, and bird species, chronic exposure to DEHP, DnBP, or DIBP has been linked to testicular toxicity (Tyler et al., 2018; Alam et al., 2025). In addition, DEHP exposure can lead to ovarian and hepatotoxicity (Alam et al., 2025). In wild populations of the fish *Squalius cephalus*, Molbert et al. (2021) found a negative association between PAE metabolite burden in muscles and telomere length, suggesting an effect on life expectancy. The concentrations measured in *E. orbicularis* in this study might thus have a deleterious impact on the general health of the individuals.

In a previous study on the same populations carried out in 2018, PCB52 and PCB153 were quantified in 45% and 41% of the individuals from Esquineau and Faïsses (Burkart et al., 2021). In the present study, we quantified PCB with low occurrence (6%), and PCB52 was detected in only 10 samples from 2019 and 2020. PCBs are persistent in the environment, and the main explanation for this discrepancy over one or two years in the same populations lies in the extraction protocol used. The method used by Burkart et al. (Burkart et al., 2021) was specifically designed to extract and purify PCB and OCP, which allowed them to obtain three to 84 times lower LOQ for these compounds than in the multiresidue method used in 2019 and 2020 (Molbert et al., 2019).

### Spatiotemporal effects

At the temporal scale, significant inter-annual variations were observed for both PAEs and PAHs, with overall higher concentrations in 2020, except for PAEs in females. These results are consistent with changes in emission patterns and meteorological conditions, particularly for PAHs (Miura et al., 2019; Dron et al., 2021). Surprisingly, plasmatic PAH burdens were higher in 2020. While lockdown periods were associated with temporary reductions in some pollutants (Bakola et al., 2022), bounce-back effects in post-lockdown periods have been observed (Li et al., 2024). We also highlighted a monthly variation for PAHs that seemed mostly driven by a few individuals with high levels in May and June (Supplementary Material, Fig. S1). Although PAEs and PAHs are continuously released by anthropogenic activities, environmental concentrations exhibit seasonal variations, with PAHs typically peaking in winter, a pattern also observed in the study area (Ravindra et al., 2008; Guigue et al., 2014). Such patterns can be reflected in biota contamination, as shown in two Mediterranean fish species (Perugini et al., 2007; Frapiccini et al., 2020), but generally require longer sampling intervals than the present study.

Globally, plasmatic burdens of PAEs and PAHs were similar between the two populations. These findings suggest that atmospheric deposition might be the primary source for these populations that are geographically close but differ in the hydrology of their habitats. For semi-volatile compounds, such as LMW PAHs and phthalates, exposure can occur via pulmonary inhalation (Teil et al. 2016), a route that is decoupled from the hydrological system and from exposure via water and freshwater prey. Although the Rhône River is a source of PAEs (Paluselli et al., 2018), turtles from the irrigation site (Esquineau) did not exhibit higher levels than the ones from the drainage site (Faïsses). We previously showed that site (and presumably the hydrology) was a significant explanatory factor in both circulating levels of MTEs and pesticides in these two populations (Merleau et al., 2024a, 2024b). In addition, PAEs and PAHs are present in substantial amounts in the atmosphere, and wet and dry depositions in freshwater environments are important factors of contamination (Ravindra et al., 2008; Bergé et al., 2013; Rabodonirina et al., 2015; Paluselli et al., 2018). At the regional scale, the ZIP of Fos-sur-Mer (Fig. 1) is a major source of PAHs, but lichen biomonitoring showed low disparity in rural and urban sampling points outside of the complex (Ratier et al., 2018). In surface waters, dissolved PAH concentrations exhibited similar concentrations in the region, except for the ZIP, with atmospheric deposition being a major source of contamination (Guigue et al., 2014). Contrasting results have been found in wildlife in the area, with high PAH disparity in eels (*Anguilla anguilla*) sampled in the Vaccares Lagoon (Ribeiro et al., 2005) but no difference in little egret (*Egretta garzetta*) eggs from three colonies of the Camargue wetland (Berny et al., 2002). All together, these results highlight that differences in the ecological characteristics of the species and in the biological matrix used lead to a contrasting pattern of contamination.

### Individual characteristics

The long-term monitoring of the two populations allowed us to assess the effect of age, which is rarely accessible in wild populations. Age was negatively correlated with PAE levels for both sexes and with PAH levels in males only. These results are consistent with the hypothesis that these compounds do not bioaccumulate in vertebrates (Mackintosh et al., 2004; Tyler et al., 2018; Goutte et al., 2020). Metabolization capacity might improve with age, leading to lower parent compounds in older individuals. Few studies have investigated the links between PAEs and age, but a positive relationship between age and PAE metabolites, but not parent compounds, has been shown in *S. cephalus* (Molbert et al., 2020). Changes in diet or habitat use have been suggested to influence PAE levels in *C. caretta* (Blasi et al., 2022) and could be linked to age in *E. orbicularis* (Ottonello et al., 2005). Growth dilution, as highlighted in *C. caretta* for PAHs and for PAE metabolites (Camacho et al., 2012; Sanjuan et al., 2023), is unlikely to play a role in *E. orbicularis*, given the absence of a link between the contaminant burden and the size of individuals in our dataset.

Sex-related differences were linked in contrasting ways to year for PAEs and age for PAHs. Higher levels of PAEs in males in 2020 only are more likely to reflect temporary changes in habitat or diet than biological differences in toxicokinetics. However, the significant decrease of PAH concentrations with age in males might indicate differential uptake/elimination or metabolization rates between sexes, as already suggested in the sea turtle *Lepidochelys kempii* (Muñoz et al., 2021). In the same populations of *E. orbicularis*, results supporting this hypothesis were observed for lead, whose levels increased in older females but not in males (Merleau et al., 2024a). In reptiles, maternal transfer has been identified for PAEs in the watersnake *Enhydris chinensis* and the sea turtle *C. caretta* (Liu et al., 2019; Savoca et al., 2021). Physico-chemical properties of the contaminants influence maternal transfer, notably the octanol-water partition coefficient (log Kow). For DnBP, DIBP, and DEHP quantified in our study, these coefficient values are comprised between 4 and 7.5 and could be compatible with a transfer to the eggs, as shown in two frog species (Liu et al., 2020). In wild species, maternal transfer of PAHs has been demonstrated in *Eretmochelys imbricata* and two fish species (Sundberg et al., 2007; Muñoz & Vermeiren, 2018). The contaminant burden quantified in *E. orbicularis* females during the reproductive season might thus have deleterious consequences for their offspring.

### Intra-individual variation across the years

The individual repeatability between years was particularly low for both PAEs and PAHs, indicating that exposure is fluctuating over time for individuals. Even though plasma is a dynamic biological compartment, these results reinforce the conclusion that PAEs and PAHs are efficiently metabolized in *E. orbicularis*, as already shown in numerous vertebrate species (Mackintosh et al., 2004; Wan et al., 2007; Fourgous et al., 2016; Tyler et al., 2018; Goutte et al., 2020). Temporal variations of PAE and PAH burdens within individuals have only been assessed in humans: urinary PAE metabolites tend to exhibit moderate to poor repeatability within individuals, but R coefficients are generally higher from 0.20 to 0.43 (Goerdten et al., 2022), than those found in *E. orbicularis* (0.065-0.108)(Goerdten et al., 2022). Further studies should also investigate plasmatic levels of PAH metabolites and phthalate monoesters to better assess exposure and toxicokinetics. Temporal variations at the intra-individual levels are contaminant-dependent: mercury, lead, and selenium had high repeatability in the same individuals (R coefficient ranging from 0.61 to 0.91), whereas bentazone, a widely used pesticide in the area, exhibited strong intra-individual variations over the years, similarly to PAEs and PAHs (Merleau et al., 2024a, 2024b). Such variability is mainly linked to the bioaccumulation potential of contaminants but also to their emission dynamics in the environment.

## Conclusion

Freshwater ecosystems constitute sinks for multiple organic contaminants, yet exposure studies remain scarce for many vertebrate species, particularly reptiles. In this study, we showed that a threatened freshwater turtle, *E. orbicularis*, presented elevated plasmatic concentrations of organic pollutants. While legacy contaminant occurrences were considerably low, PAEs and PAHs were frequently quantified at the same time. Our results suggest a local global contamination, which appears to be independent of the hydrology type of the habitats, probably originating from atmospheric deposition and fluctuating over time. Both the relationship of PAE and PAH levels with age and intra-individual variation indicate limited potential for bioaccumulation in this species, in accordance with the toxicokinetics of these compounds in vertebrates. However, the significant differences between males and females underline the need to better understand the sex-specific exposure pathways and metabolization mechanisms. In particular, monitoring the potential maternal transfer of these contaminants, which can exhibit endocrine-disrupting properties and embryotoxicity, deserves priority attention. Finally, the detection of PAEs and PAHs in these two populations, in which contamination by pesticides and MTEs has already been identified, raises additional concern for the conservation of this threatened species. Long-term monitoring efforts should be maintained to detect potential effects on individual health and population dynamics.

## Acknowledgments

The authors are grateful to Aésane Méric, Louisiane Burkart, and Louis Spinner for their help in collecting data in the field. We thank the French Ministry of Environment for giving permission to capture *E. orbicularis*.

## Data, scripts, code, and supplementary information availability

Data and script are available online: https://doi.org/10.5281/zenodo.17663455; Merleau et al., 2025.

Supplementary information is available online: https://doi.org/10.5281/zenodo.18412046; Merleau et al., 2025.

## References

Alam MS, Maowa Z, Hasan MN (2025) Phthalates toxicity in vivo to rats, mice, birds, and fish: A thematic scoping review. Heliyon, 11, e41277. 10.1016/j.heliyon.2024.e41277

Aulsebrook LC, Bertram MG, Martin JM, Aulsebrook AE, Brodin T, Evans JP, Hall MD, O’Bryan MK, Pask AJ, Tyler CR, Wong BBM (2020) Reproduction in a polluted world: implications for wildlife. Reproduction, 160, R13–R23. 10.1530/REP-20-0154

Bakola M, Hernandez Carballo I, Jelastopulu E, Stuckler D (2022) The impact of COVID-19 lockdown on air pollution in Europe and North America: a systematic review. European Journal of Public Health, 32, 962–968. 10.1093/eurpub/ckac118

Bates D, Mächler M, Bolker B, Walker S (2015) Fitting Linear Mixed-Effects Models Using lme4. Journal of Statistical Software, 67, 1–48. 10.18637/jss.v067.i01

Beau F, Bustamante P, Michaud B, Brischoux F (2019) Environmental causes and reproductive correlates of mercury contamination in European pond turtles (Emys orbicularis). Environmental Research, 172, 338–344. 10.1016/j.envres.2019.01.043

Bergé A, Cladière M, Gasperi J, Coursimault A, Tassin B, Moilleron R (2013) Meta-analysis of environmental contamination by phthalates. Environmental Science and Pollution Research, 20, 8057–8076. 10.1007/s11356-013-1982-5

Berny P, Sadoul N, Dol S, Videman B, Kayser Y, Hafner H (2002) Impact of local agricultural and industrial practices on organic contamination of little egret (Egretta garzetta) eggs in the rhône Delta, Southern France. Environmental Toxicology and Chemistry, 21, 520–526. 10.1002/etc.5620210308

Blasi MF, Avino P, Notardonato I, Di Fiore C, Mattei D, Gauger MFW, Gelippi M, Cicala D, Hochscheid S, Camedda A, De Lucia GA, Favero G (2022) Phthalate esters (PAEs) concentration pattern reflects dietary habitats (δ13C) in blood of Mediterranean loggerhead turtles (Caretta caretta). Ecotoxicology and Environmental Safety, 239, 113619. 10.1016/j.ecoenv.2022.113619

Burkart L, Olivier A, Lourdais O, Vittecoq M, Blouin-Demers G, Alliot F, Le Gac C, Martin N, Goutte A (2021) Determinants of Legacy Persistent Organic Pollutant Levels in the European Pond Turtle (Emys orbicularis) in the Camargue Wetland, France. Environmental Toxicology and Chemistry, 40, 2261–2268. 10.1002/etc.5077

Camacho M, Boada LD, Orós J, Calabuig P, Zumbado M, Luzardo OP (2012) Comparative study of polycyclic aromatic hydrocarbons (PAHs) in plasma of Eastern Atlantic juvenile and adult nesting loggerhead sea turtles (Caretta caretta). Marine Pollution Bulletin, 64, 1974–1980. 10.1016/j.marpolbul.2012.06.002

Camacho M, Luzardo OP, Boada LD, López Jurado LF, Medina M, Zumbado M, Orós J (2013) Potential adverse health effects of persistent organic pollutants on sea turtles: Evidences from a cross-sectional study on Cape Verde loggerhead sea turtles. Science of The Total Environment, 458–460, 283–289. 10.1016/j.scitotenv.2013.04.043

Campioni L, Oró-Nolla B, Granadeiro JP, Silva MC, Madeiros J, Gjerdrum C, Lacorte S (2024) Exposure of an endangered seabird species to persistent organic pollutants: Assessing levels in blood and link with reproductive parameters. Science of The Total Environment, 930, 172814. 10.1016/j.scitotenv.2024.172814

Castanet J (1988) Les méthodes d’estimation de l’âge chez les chéloniens. Mésogée, 48, 21–28.

Cocci P, Mosconi G, Bracchetti L, Nalocca JM, Frapiccini E, Marini M, Caprioli G, Sagratini G, Palermo FA (2018) Investigating the potential impact of polycyclic aromatic hydrocarbons (PAHs) and polychlorinated biphenyls (PCBs) on gene biomarker expression and global DNA methylation in loggerhead sea turtles (Caretta caretta) from the Adriatic Sea. Science of The Total Environment, 619–620, 49–57. 10.1016/j.scitotenv.2017.11.118

Dendievel A-M, Mourier B, Dabrin A, Delile H, Coynel A, Gosset A, Liber Y, Berger J-F, Bedell J-P (2020) Metal pollution trajectories and mixture risk assessed by combining dated cores and subsurface sediments along a major European river (Rhône River, France). Environment International, 144, 106032. 10.1016/j.envint.2020.106032

Dron J, Ratier A, Austruy A, Revenko G, Chaspoul F, Wafo E (2021) Effects of meteorological conditions and topography on the bioaccumulation of PAHs and metal elements by native lichen (Xanthoria parietina). Journal of Environmental Sciences, 109, 193–205. 10.1016/j.jes.2021.03.045

Dudgeon D (2019) Multiple threats imperil freshwater biodiversity in the Anthropocene. Current Biology, 29, R960–R967. 10.1016/j.cub.2019.08.002

Fay R, Ficheux S, Béchet A, Besnard A, Crochet P-A., Leblois R, Crivelli A, Wattier R, Olivier A (2023) Direct and indirect estimates of dispersal support strong juvenile philopatry and male-biased dispersal in a freshwater turtle species (Emys orbicularis). Freshwater Biology, 68, 2042–2053. 10.1111/fwb.14171

Ficheux S, Olivier A, Fay R, Crivelli A, Besnard A, Béchet A (2014) Rapid response of a long-lived species to improved water and grazing management: The case of the European pond turtle (Emys orbicularis) in the Camargue, France. Journal for Nature Conservation, 22, 342–348. 10.1016/j.jnc.2014.03.001

Fourgous C, Chevreuil M, Alliot F, Amilhat E, Faliex E, Paris-Palacios S, Teil MJ, Goutte A (2016) Phthalate metabolites in the European eel (Anguilla anguilla) from Mediterranean coastal lagoons. Science of The Total Environment, 569–570, 1053–1059. 10.1016/j.scitotenv.2016.06.159

Frapiccini E, Panfili M, Guicciardi S, Santojanni A, Marini M, Truzzi C, Annibaldi A (2020) Effects of biological factors and seasonality on the level of polycyclic aromatic hydrocarbons in red mullet (Mullus barbatus). Environmental Pollution, 258, 113742. 10.1016/j.envpol.2019.113742

Goerdten J, Yuan L, Huybrechts I, Neveu V, Nöthlings U, Ahrens W, Scalbert A, Floegel A (2022) Reproducibility of the Blood and Urine Exposome: A Systematic Literature Review and Meta-Analysis. Cancer Epidemiology, Biomarkers & Prevention, 31, 1683–1692. 10.1158/1055-9965.EPI-22-0090

González-Mille DJ, Ilizaliturri-Hernández CA, Espinosa-Reyes G, Cruz-Santiago O, Cuevas-Díaz MDC, Martín Del Campo CC, Flores-Ramírez R (2019) DNA damage in different wildlife species exposed to persistent organic pollutants (POPs) from the delta of the Coatzacoalcos river, Mexico. Ecotoxicology and Environmental Safety, 180, 403–411. 10.1016/j.ecoenv.2019.05.030

Goutte A, Alliot F, Budzinski H, Simonnet-Laprade C, Santos R, Lachaux V, Maciejewski K, Le Menach K, Labadie P (2020) Trophic Transfer of Micropollutants and Their Metabolites in an Urban Riverine Food Web. Environmental Science & Technology, 54, 8043–8050. 10.1021/acs.est.0c01411

Guigue C, Tedetti M, Ferretto N, Garcia N, Méjanelle L, Goutx M (2014) Spatial and seasonal variabilities of dissolved hydrocarbons in surface waters from the Northwestern Mediterranean Sea: Results from one year intensive sampling. Science of The Total Environment, 466–467, 650–662. 10.1016/j.scitotenv.2013.07.082

Guillot H, Bonnet X, Bustamante P, Churlaud C, Trotignon J, Brischoux F (2018) Trace Element Concentrations in European Pond Turtles (Emys orbicularis) from Brenne Natural Park, France. Bulletin of Environmental Contamination and Toxicology, 101, 300–304. 10.1007/s00128-018-2376-7

Hartig F (2024) DHARMa: Residual Diagnostics for Hierarchical (Multi-Level / Mixed) Regression Models.

Holth TF, Beylich BA, Skarphédinsdóttir H, Liewenborg B, Grung M, Hylland K (2009) Genotoxicity of Environmentally Relevant Concentrations of Water-Soluble Oil Components in Cod (Gadus morhua). Environmental Science & Technology, 43, 3329–3334. 10.1021/es803479p

Laborie S, Moreau-Guigon E, Alliot F, Desportes A, Oziol L, Chevreuil M (2016) A new analytical protocol for the determination of 62 endocrine-disrupting compounds in indoor air. Talanta, 147, 132–141. 10.1016/j.talanta.2015.09.028

Lenth RV (2025) emmeans: Estimated Marginal Means, aka Least-Squares Means.

Li R, Luo Y, Zhu X, Zhang J, Wang Z, Yang W, Li Y, Li H (2024) Anthropogenic impacts on polycyclic aromatic hydrocarbons in surface water: Evidence from the COVID-19 lockdown. Water Research, 262, 122143. 10.1016/j.watres.2024.122143

Liu Y-E, Luo X-J, Guan K-L, Huang C-C, Zhu C-Y, Qi X-M, Zeng Y-H, Mai B-X (2020) Legacy and emerging organophosphorus flame retardants and plasticizers in frogs: Sex difference and parental transfer. Environmental Pollution, 266, 115336. 10.1016/j.envpol.2020.115336

Liu Y-E, Tang B, Liu Y, Luo X-J, Mai B-X, Covaci A, Poma G (2019) Occurrence, biomagnification and maternal transfer of legacy and emerging organophosphorus flame retardants and plasticizers in water snake from an e-waste site. Environment International, 133, 105240. 10.1016/j.envint.2019.105240

Lohmann R, Breivik K, Dachs J, Muir D (2007) Global fate of POPs: Current and future research directions. Environmental Pollution, 150, 150–165. 10.1016/j.envpol.2007.06.051

López-Berenguer G, Acosta-Dacal A, Luzardo OP, Peñalver J, Martínez-López E (2023) Assessment of polycyclic aromatic hydrocarbons (PAHs) in mediterranean top marine predators stranded in SE Spain. Chemosphere, 336, 139306. 10.1016/j.chemosphere.2023.139306

Lorrain-Soligon L, Marchand E, Alliot F, Bustamante P, Petit F, Goutte A (2025) Starting from the bottom: Contrasted trophic transfer of antibiotics and pesticides through a riverine food web. Science of The Total Environment, 1002, 180617. 10.1016/j.scitotenv.2025.180617

Mackintosh CE, Maldonado J, Hongwu J, Hoover N, Chong A, Ikonomou MG, Gobas FAPC (2004) Distribution of Phthalate Esters in a Marine Aquatic Food Web: Comparison to Polychlorinated Biphenyls. Environmental Science & Technology, 38, 2011–2020. 10.1021/es034745r

Malaj E, von der Ohe PC, Grote M, Kühne R, Mondy CP, Usseglio-Polatera P, Brack W, Schäfer RB (2014) Organic chemicals jeopardize the health of freshwater ecosystems on the continental scale. Proceedings of the National Academy of Sciences, 111, 9549–9554. 10.1073/pnas.1321082111

Manzetti S (2013) Polycyclic Aromatic Hydrocarbons in the Environment: Environmental Fate and Transformation. Polycyclic Aromatic Compounds, 33, 311–330. 10.1080/10406638.2013.781042

Marasco V, Costantini D (2016) Signaling in a Polluted World: Oxidative Stress as an Overlooked Mechanism Linking Contaminants to Animal Communication. Frontiers in Ecology and Evolution, 4. 10.3389/fevo.2016.00095

Mateo R, Lacorte S, Taggart MA (2016) An Overview of Recent Trends in Wildlife Ecotoxicology. In: Current Trends in Wildlife Research Wildlife Research Monographs. (eds Mateo R, Arroyo B, Garcia JT), pp. 125–150. Springer International Publishing, Cham. 10.1007/978-3-319-27912-1_6

Matson CW, Palatnikov G, Islamzadeh A, Mcdonald TJ, Autenrieth RL, Donnelly KC, Bickham JW (2005) Chromosomal Damage in Two Species of Aquatic Turtles (Emys orbicularis and Mauremys caspica) Inhabiting Contaminated Sites in Azerbaijan. Ecotoxicology, 14, 513–525. 10.1007/s10646-005-0001-0

Matthiessen P, Wheeler JR, Weltje L (2018) A review of the evidence for endocrine disrupting effects of current-use chemicals on wildlife populations. Critical Reviews in Toxicology, 48, 195–216. 10.1080/10408444.2017.1397099

Meador JP, Stein JE, Reichert WL, Varanasi U (1995) Bioaccumulation of Polycyclic Aromatic Hydrocarbons by Marine Organisms. In: Reviews of Environmental Contamination and Toxicology Reviews of Environmental Contamination and Toxicology. (ed Ware GW), pp. 79–165. Springer New York, New York, NY. 10.1007/978-1-4612-2542-3_4

Merleau L-A, Goutte A, Olivier A, Vittecoq M, Bustamante P, Leray C, Lourdais O (2024a) Blood levels of metallic trace elements are influenced by sex, age and habitat in the European pond turtle (Emys orbicularis). Science of The Total Environment, 957, 177487. 10.1016/j.scitotenv.2024.177487

Merleau L-A, Lourdais O, Olivier A, Vittecoq M, Blouin-Demers G, Alliot F, Burkart L, Foucault Y, Leray C, Migne E, Goutte A (2024b) Pesticide concentrations in a threatened freshwater turtle (Emys orbicularis): Seasonal and annual variation in the Camargue wetland, France. Environmental Pollution, 341, 122903. 10.1016/j.envpol.2023.122903

Meyer E, Eskew EA, Chibwe L, Schrlau J, Massey Simonich SL, Todd BD (2016) Organic contaminants in western pond turtles in remote habitat in California. Chemosphere, 154, 326–334. 10.1016/j.chemosphere.2016.03.128

Miura K, Shimada K, Sugiyama T, Sato K, Takami A, Chan CK, Kim IS, Kim YP, Lin N-H, Hatakeyama S (2019) Seasonal and annual changes in PAH concentrations in a remote site in the Pacific Ocean. Scientific Reports, 9, 12591. 10.1038/s41598-019-47409-9

Molbert N, Alliot F, Goutte A, Hansson MC (2025) The dead can talk: Investigating trace element and organic pollutant exposure in mammalian roadkill under contrasting habitats. Environmental Pollution, 367, 125648. 10.1016/j.envpol.2025.125648

Molbert N, Alliot F, Leroux-Coyau M, Médoc V, Biard C, Meylan S, Jacquin L, Santos R, Goutte A (2020) Potential Benefits of Acanthocephalan Parasites for Chub Hosts in Polluted Environments. Environmental Science & Technology, 54, 5540–5549. 10.1021/acs.est.0c00177

Molbert N, Alliot F, Santos R, Chevreuil M, Mouchel J-M, Goutte A (2019) Multiresidue Methods for the Determination of Organic Micropollutants and Their Metabolites in Fish Matrices. Environmental Toxicology and Chemistry, 38, 1866–1878. 10.1002/etc.4500

Molbert N, Angelier F, Alliot F, Ribout C, Goutte A (2021) Fish from urban rivers and with high pollutant levels have shorter telomeres. Biology Letters, 17, 20200819. 10.1098/rsbl.2020.0819

Muñoz CC, Hendriks AJ, Ragas AMJ, Vermeiren P (2021) Internal and Maternal Distribution of Persistent Organic Pollutants in Sea Turtle Tissues: A Meta-Analysis. Environmental Science & Technology, 55, 10012–10024. 10.1021/acs.est.1c02845

Muñoz CC, Vermeiren P (2018) Profiles of environmental contaminants in hawksbill turtle egg yolks reflect local to distant pollution sources among nesting beaches in the Yucatán Peninsula, Mexico. Marine Environmental Research, 135, 43–54. 10.1016/j.marenvres.2018.01.012

Namroodi S, Zaccaroni A, Rezaei H, Hosseini SM (2017) European pond turtle (Emys orbicularis persica) as a biomarker of environmental pollution in Golestan and Mazandaran provinces, Iran. Veterinary Research Forum, 8, 333–339.

Net S, Sempéré R, Delmont A, Paluselli A, Ouddane B (2015) Occurrence, Fate, Behavior and Ecotoxicological State of Phthalates in Different Environmental Matrices. Environmental Science & Technology, 49, 4019–4035. 10.1021/es505233b

Oehlmann J, Schulte-Oehlmann U, Kloas W, Jagnytsch O, Lutz I, Kusk KO, Wollenberger L, Santos EM, Paull GC, Van Look KJW, Tyler CR (2009) A critical analysis of the biological impacts of plasticizers on wildlife. Philosophical Transactions of the Royal Society B: Biological Sciences, 364, 2047–2062. 10.1098/rstb.2008.0242

Olivier A (2002) Ecologie, traits d’histoire de vie et conservation d’une population de Cistude d’Europe, Emys orbicularis, en Camargue. Mémoire Ecole Pratique des Hautes Etudes.

Olivier A, Barbraud C, Rosecchi E, Germain C, Cheylan M (2010) Assessing spatial and temporal population dynamics of cryptic species: an example with the European pond turtle. Ecological Applications, 20, 993–1004. 10.1890/09-0801.1

Ottonello D, Salvidio S, Rosecchi E (2005) Feeding habits of the European pond terrapin Emys orbicularis in Camargue (Rhône delta, Southern France). Amphibia-Reptilia, 26, 562–565. 10.1163/156853805774806241

Paluselli A, Aminot Y, Galgani F, Net S, Sempéré R (2018) Occurrence of phthalate acid esters (PAEs) in the northwestern Mediterranean Sea and the Rhone River. Progress in Oceanography, 163, 221–231. 10.1016/j.pocean.2017.06.002

Perugini M, Visciano P, Giammarino A, Manera M, Di Nardo W, Amorena M (2007) Polycyclic aromatic hydrocarbons in marine organisms from the Adriatic Sea, Italy. Chemosphere, 66, 1904–1910. 10.1016/j.chemosphere.2006.07.079

Rabodonirina S, Net S, Ouddane B, Merhaby D, Dumoulin D, Popescu T, Ravelonandro P (2015) Distribution of persistent organic pollutants (PAHs, Me-PAHs, PCBs) in dissolved, particulate and sedimentary phases in freshwater systems. Environmental Pollution, 206, 38–48. 10.1016/j.envpol.2015.06.023

Rasmussen JJ, Wiberg-Larsen P, Baattrup-Pedersen A, Cedergreen N, McKnight US, Kreuger J, Jacobsen D, Kristensen EA, Friberg N (2015) The legacy of pesticide pollution: An overlooked factor in current risk assessments of freshwater systems. Water Research, 84, 25–32. 10.1016/j.watres.2015.07.021

Ratier A, Dron J, Revenko G, Austruy A, Dauphin C-E, Chaspoul F, Wafo E (2018) Characterization of atmospheric emission sources in lichen from metal and organic contaminant patterns. Environmental Science and Pollution Research, 25, 8364–8376. 10.1007/s11356-017-1173-x

Ravindra K, Sokhi R, Vangrieken R (2008) Atmospheric polycyclic aromatic hydrocarbons: Source attribution, emission factors and regulation. Atmospheric Environment, 42, 2895–2921. 10.1016/j.atmosenv.2007.12.010

Reid AJ, Carlson AK, Creed IF, Eliason EJ, Gell PA, Johnson PTJ, Kidd KA, MacCormack TJ, Olden JD, Ormerod SJ, Smol JP, Taylor WW, Tockner K, Vermaire JC, Dudgeon D, Cooke SJ (2019) Emerging threats and persistent conservation challenges for freshwater biodiversity. Biological Reviews, 94, 849–873. 10.1111/brv.12480

Relyea RA (2009) A cocktail of contaminants: how mixtures of pesticides at low concentrations affect aquatic communities. Oecologia, 159, 363–376. 10.1007/s00442-008-1213-9

Reynaud S, Deschaux P (2006) The effects of polycyclic aromatic hydrocarbons on the immune system of fish: A review. Aquatic Toxicology, 77, 229–238. 10.1016/j.aquatox.2005.10.018

Ribeiro CAO, Vollaire Y, Sanchez-Chardi A, Roche H (2005) Bioaccumulation and the effects of organochlorine pesticides, PAH and heavy metals in the Eel (Anguilla anguilla) at the Camargue Nature Reserve, France. Aquatic Toxicology, 74, 53–69. 10.1016/j.aquatox.2005.04.008

Roche H, Buet A, Jonot O, Ramade F (2000) Organochlorine residues in european eel (Anguilla anguilla), crucian carp (Carassius carassius) and catfish (Ictalurus nebulosus) from Vaccarès lagoon (French National Nature Reserve of Camargue) – effects on some physiological parameters. Aquatic Toxicology, 48, 443–459. 10.1016/S0166-445X(99)00061-2

Roche H, Vollaire Y, Martin E, Rouer C, Coulet E, Grillas P, Banas D (2009) Rice fields regulate organochlorine pesticides and PCBs in lagoons of the Nature Reserve of Camargue. Chemosphere, 75, 526–533. 10.1016/j.chemosphere.2008.12.009

Rosner B (2010) Fundamentals of Biostatistics.

Rowe CL (2008) “The Calamity of So Long Life”: Life Histories, Contaminants, and Potential Emerging Threats to Long-lived Vertebrates. BioScience, 58, 623–631. 10.1641/B580709

Sanjuan ON, Sait STL, Gonzalez SV, Tomás J, Raga JA, Asimakopoulos AG (2023) Phthalate metabolites in loggerhead marine turtles (Caretta caretta) from the Mediterranean Sea (East Spain region). Environmental Chemistry and Ecotoxicology, 5, 178–185. 10.1016/j.enceco.2023.08.003

Savoca D, Arculeo M, Barreca S, Buscemi S, Caracappa S, Gentile A, Persichetti MF, Pace A (2018) Chasing phthalates in tissues of marine turtles from the Mediterranean sea. Marine Pollution Bulletin, 127, 165–169. 10.1016/j.marpolbul.2017.11.069

Savoca D, Arculeo M, Vecchioni L, Cambera I, Visconti G, Melfi R, Arizza V, Palumbo Piccionello A, Buscemi S, Pace A (2021) Can phthalates move into the eggs of the loggerhead sea turtle Caretta caretta? The case of the nests on the Linosa Island in the Mediterranean Sea. Marine Pollution Bulletin, 168, 112395. 10.1016/j.marpolbul.2021.112395

Simms A, Robert K, Spencer R-J, Treby S, Williams-Kelly K, Sexton C, Korossy-Horwood R, Terry R, Parker A, Van Dyke J (2025) A systematic review of how endocrine-disrupting contaminants are sampled in environmental compartments: wildlife impacts are overshadowed by environmental surveillance. Environmental Science and Pollution Research, 32, 8670–8678. 10.1007/s11356-025-36211-y

Sinaei M, Zare R (2019) Polycyclic aromatic hydrocarbons (PAHs) and some biomarkers in the green sea turtles (Chelonia mydas). Marine Pollution Bulletin, 146, 336–342. 10.1016/j.marpolbul.2019.06.024

Sparling DW, Linder G, Bishop CA, Krest SK (2010) Recent Advancements in Amphibian and Reptile Ecotoxicology. In: Ecotoxicology of Amphibians and Reptiles, p.. CRC Press.

Stoffel MA, Nakagawa S, Schielzeth H (2017) rptR: repeatability estimation and variance decomposition by generalized linear mixed-effects models (S Goslee, Ed,). Methods in Ecology and Evolution, 8, 1639–1644. 10.1111/2041-210X.12797

Sundberg H, Hanson M, Liewenborg B, Zebühr Y, Broman D, Balk L (2007) Dredging Associated Effects: Maternally Transferred Pollutants and DNA Adducts in Feral Fish. Environmental Science & Technology, 41, 2972–2977. 10.1021/es070073j

Swartz CD, Donnelly KC, Islamzadeh A, Rowe GT, Rogers WJ, Palatnikov GM, Mekhtiev AA, Kasimov R, McDonald TJ, Wickliffe JK, Presley BJ, Bickham JW (2003) Chemical Contaminants and their Effects in Fish and Wildlife from the Industrial Zone of Sumgayit, Republic of Azerbaijan. Ecotoxicology, 12, 509–521. 10.1023/B:ECTX.0000003038.02643.08

Teil M-J, Moreau-Guigon E, Blanchard M, Alliot F, Gasperi J, Cladière M, Mandin C, Moukhtar S, Chevreuil M (2016) Endocrine disrupting compounds in gaseous and particulate outdoor air phases according to environmental factors. Chemosphere, 146, 94–104. 10.1016/j.chemosphere.2015.12.015

Tyler CR, Parsons A, Rogers NJ, Lange A, Brown AR (2018) Plasticisers and Their Impact on Wildlife. In: Plastics and the Environment (eds Harrison RM, Hester RE), pp. 106–130. The Royal Society of Chemistry. 10.1039/9781788013314-00106

Van Der Oost R, Beyer J, Vermeulen NPE (2003) Fish bioaccumulation and biomarkers in environmental risk assessment: a review. Environmental Toxicology and Pharmacology, 13, 57–149. 10.1016/S1382-6689(02)00126-6

Vethaak D, Legler J (2013) Endocrine Disruption in Wildlife: Background, Effects, and Implications. In: Endocrine Disrupters (ed Matthiessen P), pp. 7–58. Wiley. 10.1002/9781118355961.ch2

Wallace SJ, De Solla SR, Head JA, Hodson PV, Parrott JL, Thomas PJ, Berthiaume A, Langlois VS (2020) Polycyclic aromatic compounds (PACs) in the Canadian environment: Exposure and effects on wildlife. Environmental Pollution, 265, 114863. 10.1016/j.envpol.2020.114863

Walters DM, Jardine TD, Cade BS, Kidd KA, Muir DCG, Leipzig-Scott P (2016) Trophic Magnification of Organic Chemicals: A Global Synthesis. Environmental Science & Technology, 50, 4650–4658. 10.1021/acs.est.6b00201

Wan Y, Jin X, Hu J, Jin F (2007) Trophic Dilution of Polycyclic Aromatic Hydrocarbons (PAHs) in a Marine Food Web from Bohai Bay, North China. Environmental Science & Technology, 41, 3109–3114. 10.1021/es062594x

Yazdani M (2020) Comparative toxicity of selected PAHs in rainbow trout hepatocytes: genotoxicity, oxidative stress and cytotoxicity. Drug and Chemical Toxicology, 43, 71–78. 10.1080/01480545.2018.1497054

Yu S, Halbrook RS, Sparling DW (2012) Accumulation of Polychlorinated Biphenyls (PCBs) and Evaluation of Hematological and Immunological Effects of PCB Exposure on Turtles. Bulletin of Environmental Contamination and Toxicology, 88, 823–827. 10.1007/s00128-012-0590-2

