## Supplementary Material for "Exposure to a mixture of organic pollutants in a threatened freshwater turtle *Emys orbicularis*: effects of age, sex, and temporal variation" (from another version of this preprint: version 3, 2026-01-29)

**This file contains the revised Supplementary Material associated with the preprint "Exposure to a mixture of organic pollutants in a threatened freshwater turtle *Emys orbicularis*: effects of age, sex, and temporal variation".**

**Table S1:** Polycyclic aromatic hydrocarbons concentrations (ng.mL$^{-1}$) in plasma samples of *E. orbicularis* from the Esquineau and Faïsses populations sampled in 2019 and 2020 in the Camargue wetlands (France)

| Population | Esquineau | | | Esquineau | | | Esquineau | | | Esquineau | | |
|---|---|---|---|---|---|---|---|---|---|---|---|---|
| **Year** | **2019** | | | **2019** | | | **2020** | | | **2020** | | |
| **Sex** | **Female (N = 34)** | | | **Male (N = 24)** | | | **Female (N = 37)** | | | **Male (N = 24)** | | |
| | **Mean** | **SD** | **n** | **Mean** | **SD** | **n** | **Mean** | **SD** | **n** | **Mean** | **SD** | **n** |
| **Naphtalene** | 4.77 | 6.15 | 14 | 1.53 | 2.76 | 6 | 6.92 | 6.04 | 29 | 4.48 | 5.91 | 12 |
| **Phenanthene** | 6.25 | 33.06 | 4 | 1.17 | 2.38 | 5 | 2.24 | 5.65 | 10 | 5.61 | 13.97 | 8 |
| **Fluorene** | 1.32 | 2.90 | 11 | 0.57 | 1.14 | 5 | 0.51 | 0.99 | 8 | 0.20 | 0.71 | 2 |
| **Fluoranthene** | 0.70 | 4.08 | 1 | 0 | 0 | 0 | 0 | 0 | 0 | 8.00 | 32.32 | 2 |
| **Pyrene** | 0.43 | 2.49 | 1 | 0 | 0 | 0 | 0 | 0 | 0 | 10.22 | 44.53 | 2 |
| **Dibenzo_a_h_anthracene** | 1.09 | 3.90 | 3 | 0 | 0 | 0 | 0 | 0 | 0 | 0 | 0 | 0 |
| **Indeno_123_cd_pyrene** | 0.44 | 2.54 | 1 | 0 | 0 | 0 | 0 | 0 | 0 | 0 | 0 | 0 |
| **Benzo_g_h_i_perylene** | 0.26 | 1.49 | 1 | 0 | 0 | 0 | 0 | 0 | 0 | 0 | 0 | 0 |
| **Population** | **Faisses** | | | **Faisses** | | | **Faisses** | | | **Faisses** | | |
| **Year** | **2019** | | | **2019** | | | **2020** | | | **2020** | | |
| **Sex** | **Female (N = 29)** | | | **Male (N = 20)** | | | **Female (N = 34)** | | | **Male (N = 16)** | | |
| | **Mean** | **SD** | **n** | **Mean** | **SD** | **n** | **Mean** | **SD** | **n** | **Mean** | **SD** | **n** |
| **Naphtalene** | 4.59 | 6.11 | 14 | 3.75 | 5.52 | 7 | 6.26 | 5.64 | 24 | 9.70 | 8.22 | 12 |
| **Phenanthene** | 0.76 | 1.93 | 4 | 2.77 | 6.53 | 6 | 4.98 | 12.46 | 12 | 2.32 | 6.22 | 3 |
| **Fluorene** | 0.91 | 1.42 | 9 | 1.19 | 1.77 | 7 | 0.17 | 0.57 | 3 | 0.89 | 1.51 | 5 |
| **Fluoranthene** | 0.12 | 0.67 | 1 | 0 | 0 | 0 | 0 | 0 | 0 | 0 | 0 | 0 |
| **Pyrene** | 0 | 0 | 0 | 0 | 0 | 0 | 0 | 0 | 0 | 0 | 0 | 0 |
| **Dibenzo_a_h_anthracene** | 0 | 0 | 0 | 0 | 0 | 0 | 0 | 0 | 0 | 0 | 0 | 0 |
| **Indeno_123_cd_pyrene** | 0 | 0 | 0 | 0 | 0 | 0 | 0 | 0 | 0 | 0 | 0 | 0 |
| **Benzo_g_h_i_perylene** | 0 | 0 | 0 | 0 | 0 | 0 | 0 | 0 | 0 | 0 | 0 | 0 |

N: number of individuals sampled (by site, year, and sex)

SD: standard deviation

n: number of samples with concentrations > limits of quantification

**Table S2:** Phthalate diester concentrations (ng.mL$^{-1}$) in plasma samples of *E. orbicularis* from the Esquineau and Faïsses populations sampled in 2019 and 2020 in the Camargue wetlands (France)

| **Population** | **Esquineau** | | | **Esquineau** | | | **Esquineau** | | | **Esquineau** | | |
|---|---|---|---|---|---|---|---|---|---|---|---|---|
| **Year** | **2019** | | | **2019** | | | **2020** | | | **2020** | | |
| **Sex** | **Female (N =34)** | | | **Male (N = 24)** | | | **Female (N = 37)** | | | **Male (N = 24)** | | |
| | **Mean** | **SD** | **n** | **Mean** | **SD** | **n** | **Mean** | **SD** | **n** | **Mean** | **SD** | **n** |
| **DMP** | 3.79 | 5.4 | 13 | 4.54 | 9.2 | 7 | 1.27 | 2.9 | 7 | 3.58 | 6.6 | 8 |
| **DEP** | 27.44 | 93.7 | 4 | 43.13 | 91.7 | 6 | 26.03 | 75.9 | 7 | 102.54 | 176.4 | 10 |
| **DIBP** | 142.38 | 372.3 | 10 | 146.83 | 265.0 | 7 | 298.30 | 667.2 | 22 | 958.79 | 1223.8 | 20 |
| **DNBP** | 92.85 | 260.9 | 11 | 131.71 | 252.1 | 7 | 150.11 | 338.1 | 13 | 504.71 | 638.0 | 18 |
| **DEHP** | 9.21 | 16.9 | 12 | 40.25 | 100.5 | 18 | 21.08 | 67.4 | 9 | 123.92 | 510.3 | 11 |
| **Population** | **Faisses** | | | **Faisses** | | | **Faisses** | | | **Faisses** | | |
| **Year** | **2019** | | | **2019** | | | **2020** | | | **2020** | | |
| **Sex** | **Female (N = 29)** | | | **Male (N = 20)** | | | **Female (N = 34)** | | | **Male (N = 16)** | | |
| | **Mean** | **SD** | **n** | **Mean** | **SD** | **n** | **Mean** | **SD** | **n** | **Mean** | **SD** | **n** |
| **DMP** | 11.38 | 21.7 | 16 | 7.25 | 12.7 | 6 | 2.79 | 6.7 | 8 | 6.31 | 12.1 | 7 |
| **DEP** | 26.59 | 66.4 | 5 | 23.75 | 64.1 | 3 | 28.85 | 72.8 | 7 | 87.31 | 209.3 | 5 |
| **DIBP** | 226.48 | 489.5 | 8 | 291.40 | 1014.2 | 5 | 359.24 | 715.6 | 21 | 1001.19 | 1905.3 | 14 |
| **DNBP** | 92.76 | 206.1 | 9 | 77.70 | 227.2 | 5 | 195.76 | 391.0 | 19 | 373.50 | 751.7 | 12 |
| **DEHP** | 22.55 | 83.4 | 12 | 22.75 | 54.3 | 8 | 18.68 | 37.0 | 15 | 20.56 | 45.2 | 6 |

N: number of individuals sampled (by site, year, and sex)

SD: standard deviation

n: number of samples with concentrations > to limits of quantification

**Table S3:** Internal standard solutions and volumes used for internal calibration of the plasma samples

| Compounds | Initial Concentration (ng/µL) | Volume (µL) | Final Quantity (ng) |
|---|---|---|---|
| Phenanthrene d10, Pyrene d10, Benzo(a)anthracene d12 (Cluzeau Info Labo) | 3.33 | 10 | 33.3 |
| PAEs: DEP-d4 and DEHP d4 (LGC Starndards) | 10 | 20 | 200 |
| Lindage d6 (LGC Starndards) | 10 | 5 | 50 |
| Cis-Permethrine 13C (LGC Starndards) | 10 | 10 | 100 |
| PCB 30, PCB 107 (Cluzeau Info Labo) | 1 | 10 | 10 |
| $^{13}C_{12}$-BDE-47, $^{13}C_{12}$-BDE153 (BCP Instruments) | 0.494 | 10 | 4.94 |
| Acetone | - | 50 | - |

**Table S4**: Selected GC–MS/MS parameters for parent compounds and their internal standards (RT: retention time, Q: quantifier transition and q: qualifier transition)

| Compound | RT (min) | Quantification transition (Q) | Q Collision energy (eV) | Confirmation transition(q) | q Collision energy (eV) | q/Q ratio (%) |
|---|---|---|---|---|---|---|
| Naphtalene | 7.4 | 128 >102 | 20 | 128 > 77 | 45 | 19.7 |
| DMP | 9.97 | 163 > 77 | 25 | 163 > 133 | 10 | 25.8 |
| Acenaphtylene | 10.12 | 152 > 126 | 25 | 152 > 150 | 35 | 97.9 |
| Acenaphtene | 10.6 | 154 > 153 | 15 | 154 > 152 | 35 | 41.4 |
| PeCB * | 10.9 | 250 > 215 | 25 | 250 > 142 | 40 | 64.3 |
| DEET * | 11.95 | 119 > 91 | 10 | 119 > 65 | 30 | 41.6 |
| Fluorene | 12.15 | 166 > 165 | 20 | 166 > 163 | 55 | 7.5 |
| DEP | 12.05 | 149 > 65 | 25 | 149 > 121 | 10 | 52.9 |
| DEP-$d_4$ * | 12.1 | 153 > 97 | 10 | 153 > 125 | 10 | 52.9 |
| HCB * | 14.2 | 249 > 214 | 13 | 284 > 249 | 18 | 24.7 |
| PCB 30 | 15.3 | 256 > 186 | 25 | 256 > 151 | 25 | - |
| Lindane-$d_6$ * | 15.3 | 224 > 187 | 10 | 224 > 150 | 20 | 96.6 |
| Lindane | 15.6 | 216.8 > 180.8 | 10 | 180.8 > 145 | 20 | 96.6 |
| Phenanthrene-$d_{10}$ | 16.1 | 188 > 184 | 45 | 188 > 160 | 45 | 84.3 |
| Phenanthrene | 16.25 | 178 > 152 | 30 | 178 > 176 | 45 | 39.3 |
| Anthracene | 16.4 | 178 > 176 | 50 | 178 > 152 | 25 | 46.7 |
| DiBP * | 17.95 | 149 > 121 | 20 | 223 > 149 | 20 | 9.1 |
| PCB 28 | 18.2 | 256 > 186 | 25 | 256 > 151 | 25 | 24.1 |
| PCB 52 | 20 | 292 > 222 | 27 | 292 > 220 | 27 | 73.2 |
| DnBP * | 20.4 | 149 > 65 | 30 | 149 > 93 | 20 | 24.9 |
| Fluoranthene | 23.05 | 202 > 200 | 45 | 202 > 150 | 55 | 19.5 |
| Pyrene-$d_{10}$ | 24.3 | 212 > 208 | 45 | 212 > 210 | 30 | 84.2 |
| Pyrene | 24.4 | 202 > 200 | 45 | 202 > 150 | 55 | 17.7 |
| PCB 101 | 27.9 | 326 > 256 | 27 | 326 > 254 | 27 | 82.3 |
| p,p'-DDE | 25.2 | 246 > 176 | 20 | 318 > 248 | 20 | 47.6 |
| PCB 107 * | 27.6 | 326 > 256 | 27 | 326 > 254 | 27 | - |
| BDE 28 | 28.2 | 407.8 > 247.9 | 20 | 247.9 > 139 | 30 | 1.7 |
| PCB 153 | 29 | 360 > 290 | 27 | 360 > 325 | 12 | 47.8 |
| PCB 138 | 30.5 | 360 > 290 | 27 | 360 > 325 | 12 | 33.3 |
| BBP | 30.7 | 149 > 65 | 25 | 149 > 121 | 10 | 51.4 |
| Benzo(a)anthracene | 33 | 228 > 226 | 45 | 228 > 224 | 55 | 34.5 |
| Benzo(a)anthracene-$d_{12}$ * | 32.9 | 240 > 236 | 40 | 240 > 238 | 15 | 34.5 |
| Chrysene | 33.3 | 228 > 226 | 45 | 228 > 224 | 55 | 28.8 |
| Bifenthrine * | 33.8 | 181 > 166 | 25 | 181 > 165 | 20 | 37.7 |
| PCB 118 | 34.3 | 326 > 256 | 27 | 326 > 254 | 27 | 85.9 |
| PCB 180 | 34.2 | 394 > 324 | 20 | 394 > 359 | 20 | 33.3 |
| BDE 47 | 34.6 | 485.7 > 325.9 | 25 | 325.9 > 138 | 55 | 2.8 |
| $^{13}$C-PBDE 47 * | 35 | 497.7 > 337.9 | 25 | 337.9 > 148.9 | 55 | 2.8 |
| Permethrine * | 35 | 183.1 > 153 | 15 | 183,1 > 168 | 15 | 73.6 |
| Phenothrine * | 35.4 | 183.1 > 153 | 15 | 183,1 > 168 | 15 | 68.7 |
| DEHP | 35.3 | 149 > 65 | 25 | 149 > 121 | 10 | 50.7 |
| DEHP-$d_4$ * | 35.3 | 153 > 125 | 10 | 153 > 97 | 10 | 50.7 |
| BDE 100 | 38.5 | 403.7 > 137 | 55 | 565.7 > 405.9 | 25 | 71.1 |
| DnOP * | 39 | 149 > 65 | 25 | 149 > 93 | 20 | 35.4 |
| Benzo(b)fluoranthene | 39.2 | 252 > 250 | 45 | 250 > 248 | 50 | 10.0 |
| Benzo(k)fluoranthene | 39.35 | 252 > 250 | 45 | 250 > 248 | 45 | 9.6 |
| BDE 99 | 39.5 | 403.7 > 137 | 55 | 565.7 > 405.9 | 25 | 47.5 |
| Cyfluthrine * | 39.6-40.2 | 163 > 127 | 10 | 226 > 206 | 10 | 26.3 |
| Benzo(a)pyrene | 40.6 | 252 > 250 | 45 | 252 > 224 | 55 | 13.1 |
| $^{13}$C-Permethrine * | 40.4 | 189 > 174 | 10 | 189 > 161 | 5 | 73.6 |
| Cypermethrine * | 40.6-41 | 163 > 127 | 5 | 181 >152 | 20 | 46.4 |
| Fenvarelate * | 42.2-42.6 | 167 > 125 | 10 | 225 > 119 | 15 | 23.0 |
| BDE_154 | 42 | 643.6 > 483.9 | 25 | 483.9 > 376.8 | 40 | 82.8 |
| BDE_153 | 43.5 | 643.6 > 483.9 | 25 | 483.9 > 376.8 | 40 | 46.8 |
| $^{13}$C-PBDE 153 * | 44 | 655.7 > 495.8 | 25 | 495.7 > 335.8 | 40 | 46.8 |
| Deltamethrine * | 43.4-43.8 | 253 > 172 | 10 | 181 > 152 | 10 | 68.9 |
| Indeno(1,2,3-cd)pyrene | 45.1 | 276 > 274 | 45 | 276 > 272 | 55 | 10.2 |
| Dibenzo(a,h)anthracene | 45.3 | 278 > 276 | 40 | 278 > 274 | 55 | 24.3 |
| Benzo(g,h,i)perylene | 45.7 | 276 > 274 | 50 | 276 > 272 | 55 | 9.1 |

* MRM conditions optimized

**Table S5**: Recovery yields, standard deviations, limits of detection and quantification, and instrumental detection limits of the compounds from the seven chemical families

| Chemical Family/Compound | % RR | % RSD | LOQ ng/mL | LOD ng/mL | IDL (pg inj.) |
|---|---|---|---|---|---|
| **Phthalates** | | | | | |
| DMP | 82 | 1.6 | 3.9 | 1.3 | 0.01 |
| DEP | 66 | 2.2 | 49.1 | 16.4 | 0.06 |
| DIBP | 83 | 15.3 | 65.9 | 22.0 | 0.04 |
| DNBP | 220 | 3.4 | 54 | 18.0 | 0.28 |
| BBP | 103 | 4.5 | 28.4 | 9.5 | 0.66 |
| DEHP | 84 | 1 | 4.9 | 1.6 | 0.02 |
| DnOP | 54 | 5.6 | 7.2 | 2.4 | 0.22 |
| **Polycylic Aromatic Hydrocarbons** | | | | | |
| Acenaphthene | 75 | 3.7 | 15.9 | 5.3 | 0.05 |
| Acenaphthylene | 56 | 5.9 | 10 | 3.3 | 0.04 |
| Naphthalene | 70 | 4.2 | 4.7 | 1.6 | 0.03 |
| Anthracene | 39 | 6.1 | 5.9 | 2.0 | 0.13 |
| Fluorene | 78 | 4.1 | 1.8 | 0.6 | 0.05 |
| Phenanthrene | 129 | 0.8 | 4.1 | 1.4 | 0.11 |
| Fluoranthene | 43 | 2.6 | 3.6 | 1.2 | 0.29 |
| Pyrene | 169 | 1.4 | 8.4 | 2.8 | 0.23 |
| Benzo_a_anthracene | 154 | 3.3 | 5.5 | 1.8 | 0.24 |
| Chrysene | 41 | 2.8 | 3.6 | 1.2 | 0.16 |
| Benzo_a_pyrene | 37 | 11.1 | 10.2 | 3.4 | 0.66 |
| Benzo_b_fluoranthene | 44 | 4.1 | 8.7 | 2.9 | 0.09 |
| Benzo_k_fluoranthene | 45 | 6.2 | 6 | 2.0 | 0.08 |
| Benzo_g_h_i_perylene | 42 | 10.2 | 7.7 | 2.6 | 0.08 |
| Dibenzo_a h_anthracene | 88 | 2.3 | 5.7 | 1.9 | 0.1 |
| Indeno_1 2 3-cd_pyrene | 149 | 2.3 | 14.2 | 4.7 | 0.28 |
| **Pyrethrinoides** | | | | | |
| Bifenthrin | 181 | 8.1 | 15.0 | 5.0 | 1.02 |
| Permethrin | 192 | 6.8 | 15.0 | 5.0 | 11.25 |
| Phenothrin | 138 | 5.2 | 15.0 | 5.0 | 10.23 |
| Cyfluthrin | 154 | 17.6 | 15.0 | 5.0 | 11.84 |
| Cypermethrin | 120 | 19.1 | 15.0 | 5.0 | 13.2 |
| Fenvarelate | 175 | 21.2 | 15.0 | 5.0 | 4.22 |
| Deltamethrin | 181 | 16.8 | 15.0 | 5.0 | 6.43 |
| **DEET** | 131 | 11.1 | 33.8 | 11.3 | 13.64 |
| **Organochlorine Pesticides** | | | | | |
| PeCB | 99 | 10.2 | 4.9 | 1.6 | 0.02 |
| HCB | 104 | 9.6 | 16.9 | 5.6 | 0.06 |
| Lindane | 106 | 7 | 22.5 | 7.5 | 0.11 |
| 44'-DDE | 115 | 6.7 | 3.6 | 1.2 | 0.02 |
| **PCB** | | | | | |
| 28 | 127 | 2 | 1.7 | 0.6 | 0.01 |
| 52 | 108 | 1.6 | 3.1 | 1.0 | 0.03 |
| 101 | 118 | 3.3 | 2 | 0.7 | 0.03 |
| 138 | 91 | 3.5 | 2 | 0.7 | 0.08 |
| 153 | 92 | 3 | 2.2 | 0.7 | 0.06 |
| 118 | 89 | 5.1 | 9.6 | 3.2 | 0.19 |
| 180 | 89 | 2.9 | 4.2 | 1.4 | 0.11 |

| PBDE | % RR | % RSD | LOQ ng/mL | LOD ng/mL | IDL (pg inj.) |
|---|---|---|---|---|---|
| 28 | 99 | 7.6 | 3.9 | 1.3 | 0.05 |
| 47 | 95 | 5.2 | 1.1 | 0.4 | 0.13 |
| 100 | 65 | 4 | 11 | 3.7 | 0.32 |
| 99 | 68 | 2.6 | 11 | 3.7 | 0.32 |
| 154 | 59 | 2.8 | 4.7 | 1.6 | 0.54 |
| 153 | 64 | 8.2 | 4.5 | 1.5 | 0.87 |

RR: recovery rate; RSD: relative standard deviation; LOQ: limit of quantification; LOD: limit of detection; IDL: instrumental detection limit
PeCB: pentachlorobenzene; HCB: hexachlorobenzène; 44'-DDE:

**Fig S1:** Sum (Σ) of the three polycyclic aromatic hydrocarbons (PAH) concentrations ($ng.mL^{-1}$) quantified in the plasma of *E. orbicularis* from two populations in the Camargue (France) sampled in 2019 and 2020, according to the month of sampling.

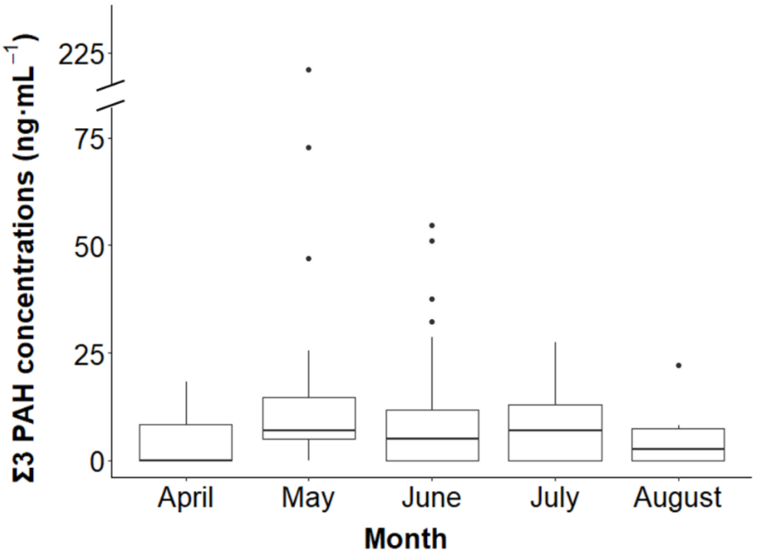
